# Overwintering under the ice: seasonal shifts in cold and hypoxia stress physiology of three winter-active pond insects

**DOI:** 10.64898/2026.08.05.743153

**Authors:** Luke S. Burton, Teresa A. Keenan, Tamara M. Rodela, Jantina Toxopeus

**Author notes:** corresponding author: JT. **Author emails**: LSB; TAK; TMR.

## Abstract

Winter poses harsh physiological challenges for insects living in temperate ponds due to the combination of low temperatures and reduced oxygen levels (hypoxia). Despite these stressors, water boatmen (*Hesperocorixa* sp.), backswimmers (*Notonecta* sp.) and diving beetles (*Laccophilus maculosus*) remain active year-round in eastern Nova Scotia, including in ice-covered ponds. However, the underlying mechanisms that allow pond insects to survive overwintering have not been widely studied. We hypothesized that cold and hypoxia tolerance would improve from September to March in these insects, with changes in whole-animal and biochemical correlates of tolerance to both stressors. We collected insects from the field and stocked them in outdoor freshwater mesocosms. Every two months, between September 2023 and April 2024, we characterized whole animal responses and biochemical changes. Whole insect cold tolerance assays indicated a trend of improved ability to sustain voluntary muscle control (CT_min_) at lower temperatures in winter-collected insects and higher internal fluid freezing temperatures (SCP). There were no changes in whole animal correlates of hypoxia tolerance over time (surface respiration frequency, submersion and surface time). Potential cryoprotectants like proline, *myo*-inositol and trehalose increased in concentration in multiple insect taxa during winter but were relatively low compared to terrestrial insect studies. As lactate dehydrogenase activity did not change, there is little evidence that the insects experienced functional hypoxia sufficient to induce anaerobic metabolism. Overall, this research has established a baseline cold and hypoxia tolerance dataset in understudied pond insects.

**Summary Statement:** Pond insects remain active during low temperature and low oxygen conditions associated with ice cover in winter, supported by physiological and biochemical changes.

## Introduction

Many organisms are exposed to a variety of co-occurring environmental stressors in their natural habitats. The ability of animals to tolerate stressors depends on the species-specific physiological responses and the spatial and temporal scales of the stressors themselves (Kaunisto et al., 2016). The physiological mechanisms of multiple stress tolerance can interact positively or negatively with one another. Some organisms are cross-tolerant to multiple stressors: the acclimatization or tolerance to one stressor can aid in the acclimatization to another stressor, particularly if common physiological tolerance mechanisms are involved (Pallarés et al., 2017; Sinclair et al., 2013). For example, aquatic beetles effectively protect against two stressors – salinity and desiccation – using a common mechanism: regulation of water balance (Pallarés et al., 2017). Conversely, when an organism is cross-susceptible to multiple stressors, the acclimatization to one stressor can be energetically costly, and deplete the physiological resources needed to contend with other stressor(s) (Krebs and Feder, 1998; Todgham and Stillman, 2013). Two commonly studied co-occurring stressors in temperate, freshwater ponds are high temperature and low oxygen (Verberk and Bilton, 2013), but few studies have explored the effects of low temperature and hypoxia in these environments.

Low temperature and hypoxia likely co-occur in ice-covered ponds with limited water inflow or outflow during winter, potentially causing challenges for overwintering aquatic insects. In temperate zones, water temperatures decrease in the late fall and typically remain between 0 and 4°C throughout winter (Brittain and Nagell, 1981; Danks, 2008; Lauff et al., 2025).

Following consistent subzero air temperatures in early winter (e.g., December or January in northern climates), seasonal ice and snow coverage can form a physical barrier that restricts atmospheric gas exchange and simultaneously reduce photosynthetic oxygenation, resulting in depletion of dissolved oxygen (DO) during the winter (Jansen et al., 2025; Mathias and Barica, 1980; Zhang et al., 2024). Despite this seasonal dynamic of cold and hypoxia stressors, families of insects from more than six orders have been sampled alive and motile during the winter, underneath more than 10 cm of pond ice in eastern Nova Scotia, Canada (c. 45°N; Lauff et al., 2025). In our study, we characterized the effect of low temperature and hypoxia on overwintering, active pond insects.

We focused on three species of winter-active air-breathing pond insects: backswimmers (*Notonecta* sp.), water boatmen (*Hesperocorixa* sp.), and diving beetles (*Laccophilis maculosus*). Backswimmers and water boatmen are morphologically similar Hemiptera (true bugs) that are widely dispersed (Wang et al., 2021). *Notonecta* and *Hesperocorixa* are common genera of backswimmers and water boatmen, respectively, each with multiple species found in Nova Scotia (Table S1). Dytiscidae is a large family of aquatic Coleoptera (beetles; Jones and Seymour, 2021; White and Roughley, 2008). Among them, *Laccophilus* is a genus of relatively small diving beetles that is abundant in ponds throughout Eastern Nova Scotia (Lauff et al., 2025). In Nova Scotia, only a single species, *Laccophilus macuolosus,* has been reported (Table S1).

Like their terrestrial relatives, adult water bugs and beetles rely on their tracheal system for gas exchange, but are bimodal breathers that can extract oxygen from both the atmosphere and water (Seymour and Matthews, 2013). Backswimmers and boatmen use a plastron, whereas, diving beetles use an air bubble, both of which trap air from the water surface on their bodies and remain connected to the tracheal system underwater (Wigglesworth, 1930). In addition to serving as an oxygen reservoir, these air stores also function as a “physical gill” (rather than anatomical gill; Seymour and Matthews, 2013). The physical gill is a gas permeable surface that enables dissolved oxygen to diffuse continuously from the surrounding water into the bubble (Seymour and Matthews, 2013). Oxygen uptake is driven by the partial pressure (PO_2_) gradient between the surrounding water and the air bubble, partially replenishing oxygen consumed during extended dive durations (Seymour and Matthews, 2013; Seymour et al., 2015). Consequently, respiratory gas exchange is closely linked to diving behaviours, including dive duration, surfacing frequency, and time spent at the water surface (Jones and Seymour, 2021). As environmental oxygen declines, the effectiveness of the physical gill decreases, making diving behaviour a key determinant of tissue oxygenation.

Functional hypoxia occurs when oxygen delivery to the tissues becomes insufficient to maintain normal aerobic metabolism and physiological function (Harrison et al., 2018). Under these condition, aquatic air-breathing insects are expected to increase visits to the water surface (aquatic surface respiration; ASR) to to replenish their air stores with atmospheric oxygen (Carbonell et al., 2024; De Ruiter et al., 1951; Gittelman, 1975). If access to the water surface is prevented by sustained winter ice cover, or if insects cannot only extract oxygen from the water (e.g., via anatomical or physical gills), insects may become increasingly reliant on anerobic metabolism (Srayko et al., 2023). Although anaerobic glycolysis can temporarily sustain ATP production, it yields substantially less ATP than aerobic respiration, and can deplete of energy reserves (e.g., glycogen) while causing an accumulation of metabolic by-products such as lactate (Farhana and Lappin, 2022; Sokolova, 2013). These energetic constraints are likely to be important for winter-active insects, which must maintain locomotor activity throughout the winter.

Insects can respond to hypoxic conditions through behavioural and physiological adjustments (Frakes et al., 2021). To behaviourally avoid hypoxic pond environments, insects may migrate to more oxygen-rich habitats such as lakes or rivers in the fall (Danks, 2008; Srayko et al., 2022). To decrease metabolic demand for oxygen in hypoxic environments, motile animals can limit their activity (Richards, 2009). Air-breathing insects that cannot access surface oxygen may use their physical gill (air bubble or plastron) to obtain dissolved oxygen from the surrounding water, but only if there is a sufficient gradient of oxygen (Carbonell et al., 2023; Gittelman, 1975). Aquatic gill-breathing mayfly larvae can use anaerobic metabolism to subsidize ATP demands under short (1 h) hypoxic conditions, as indicated by increased lactate dehydrogenase (LDH) activity (Brittain and Nagell, 1981; Cochran et al., 2022; Verberk et al., 2013), but prolonged anaerobic metabolism can be harmful. The mechanisms of hypoxia stress and tolerance in air-breathing aquatic insects, such as coleopterans and hemipterans, are understudied compared with those in gill-breathing or terrestrial insects, particularly in combination with low temperatures.

Exposure to temperatures near 0°C during winter months in temperate ponds can present an array of physiological challenges. Mild low temperatures can severely hinder protein function (e.g., enzyme activity) and decrease cellular membrane fluidity (Overgaard and MacMillan, 2017; Somero et al., 2017). This can impair ATP production and maintenance of ion balance, which can lead to loss of neuromuscular function and whole-animal bodily control. The low temperature at which bodily control is lost is referred to as the critical thermal minimum (CT_min_) (Denlinger and Lee, 2010; MacMillan et al., 2012). Prolonged exposure to mild low temperatures can also cause chilling injury (Overgaard and MacMillan, 2017). While there is no risk of freezing (internal ice formation) in water temperatures between 0 to 4 °C, aquatic insects can be encapsulated within winter ice and may freeze due to contact with external ice at temperatures below 0°C (Danks, 2008; Srayko et al., 2023; Toxopeus and Sinclair, 2018). Internal ice formation occurs at a subzero temperature, referred to as the supercooling point (SCP), and is lethal for most species of insects (Sinclair et al., 2015; Toxopeus and Sinclair, 2018). Thus, overwintering pond conditions can limit insect activity, cause chilling injury, and potentially induce lethal freezing of insects – especially those in contact with ice.

To protect against low temperature stress, insects can alter their behaviour and physiology. Aquatic pond insects may behaviourally avoid cold exposure by overwintering in different microhabitats in the same or new body of water. For example, various dipteran (fly) and coleopteran (beetle) species overwinter in the sediment or under leaf litter (Lencioni, 2004; Mihalicz, 2015). If changes in behaviour cannot sufficiently mitigate cold stress, winter survival depends on physiological mechanisms of cold hardiness, most of which have been described in terrestrial insects (Overgaard and MacMillan, 2017; Renault et al., 2002). To stay active at low temperatures (i.e., lower the CT_min_), insects need to maintain ion balance, which is critical for muscular coordination and general bodily functions (Overgaard and MacMillan, 2017). Insects can alter cell membrane composition to maintain fluidity at lower temperatures, which aids in ion homeostasis and selective permeability (Koštal et al., 2004; Overgaard and MacMillan, 2017; Tomcala et al., 2006). To prevent freezing in subzero temperatures, insects can depress (decrease) their SCP by accumulating antifreeze proteins (AFPs) and low molecular weight cryoprotectants such as sugars, polyols and amino acids (Hawes et al., 2011; Steele, 1982; Wan et al., 2023). Glycogen is a primary substrate from which various metabolites (e.g., glucose, glycerol and trehalose) can be derived from to function as cryoprotectants or metabolic fuel (Hochachka et al., 2022; Pichaud et al., 2025; Wan et al., 2023). Cryoprotectants colligatively depress the SCP by increasing internal solute concentration without harmful effects, as has been observed in insects like emerald ash borers, lady beetles, and goldenrod gall moths (Crosthwaite et al., 2011; Kelleher et al., 1987; Watanabe, 2002). Cryoprotectants can also protect cellular structures (e.g., membranes, proteins) via non-colligative mechanisms (Toxopeus et al., 2019a). The extent to which aquatic insects use the cold tolerance mechanisms described in terrestrial insects is not well-studied.

In this study, we hypothesized that both cold and hypoxia tolerance would improve prior to or during the winter to facilitate the survival and activity of overwintering water boatmen, backswimmers, and diving beetles. In addition, we hypothesized that cold tolerance would increase before hypoxia tolerance because temperatures decrease in the fall, while oxygen is only expected to decrease after ice cover is sustained in early winter. With respect to cold tolerance, we predicted that overwintering insects would sustain motility at lower temperatures (decreased CT_min_), accumulate low molecular weight cryoprotectants, and exhibit lower SCPs compared to fall and spring insects. We predicted that improved hypoxia tolerance following sustained ice cover would correlate with improved dive capacity (lower ASR frequency and more time spent submerged) and the ability to use anaerobic metabolism (increased LDH activity and depleted glycogen) during short hypoxia exposures.

## Methods

### Study species and genetic identification

Notonectidae (backswimmers), Corixidae (water boatmen), and Dytiscidae (diving beetles) were selected for their abundance and ecological relevance in ponds in Nova Scotia, Canada. We morphologically and genetically identified the three taxa used in our study. Insects were morphologically identified to family and genus using taxonomic dichotomous keys (White and Roughley, 2008; Polhemus, 2008). For genetic identification, we first extracted DNA from the legs of 4 individuals per insect taxon using a Pure Gene DNA Extraction kit (Qiagen, Toronto, Ontario, Canada) according to manufacturer’s instructions (Lemay et al., 2024). We amplified a 3’ portion of the *Cytochrome c Oxidase Subunit 1 (COI)* gene using TL2-N-3014 (Pat) and C1-J-2183 (Jerry) primers (Simon et al., 1994) and Phusion High-Fidelity Taq polymerase (Thermo Fisher, Waltham, Massachusetts, USA) according to the manufacturer’s instructions. PCR conditions included an initial denaturation at 98°C for 30 s; followed by 40 cycles of denaturation at 98°C for 10 s, annealing at 52°C for 30 s, extension at 72°C for 30 s; and a final extension at 72°C for 5 min. We confirmed amplicon sizes on a 1.5% agarose gel and sent the PCR products to The Centre for Applied Genomics (Sick Kids Hospital, Toronto, Ontario, Canada) for Sanger sequencing. *COI* sequences were compared to other *COI* sequences deposited in NCBI (https://www.ncbi.nlm.nih.gov/) using BLAST (basic local alignment search tool). Clustal Omega (Madeira et al., 2024; https://www.ebi.ac.uk/jdispatcher/msa/clustalo) was used to compare sequences within each taxon to each other. Based on BLAST results, water boatmen (*Hesperocorixa* sp.) and backswimmers (*Notonecta* sp.) were identified to the genus level, and diving beetles (*Laccophilus maculosus*) to the species level (Table S1). Based on Clustal Omega analyses, there was high similarity among genetic sequences within each genus we sampled (Fig. S1), suggesting that we captured only one species each of the water boatmen, backswimmers, and diving beetles.

### Experimental design and insect collection

To determine if low temperature and hypoxia tolerance increased during winter in our three target taxa, we measured whole animal and biochemical indicators of stress tolerance in insects acclimatized to semi-natural (mesocosm) or natural (pond) conditions at four seasonal time points, which included end of summer (late September to mid-October, 2023), end of autumn/early winter (late November to mid-December, 2023), mid-winter (late January to mid-February, 2024) and end of winter/early spring (late March to mid-April, 2024). Mesocosms were stocked with field-collected insects prior to the start of the experiment, and these stocks were sufficiently high to measure correlates of stress tolerance as planned between September and November but insect abundance declined in January and were near-zero by March (Table 1). There were insufficient diving beetles to complete most measurements for the January time point, except for whole animal hypoxia tolerance. All insects used in March experiments were collected from the two field sites that the tanks were originally stocked from. At each time point, we brought animals into the laboratory to measure CT_min_ (in water) and SCP (in air) as metrics of whole-animal cold tolerance. In separate animals, we measured hypoxia tolerance by observing surface respiration frequency and duration when exposed to hypoxic mesocosm or pond water in the laboratory. We also measured concentrations of low molecular weight cryoprotectants, glycogen, and LDH activity in whole body homogenates of each taxon at each time point.

**Table 1.** Number of individuals from each taxon present in mesocosms before the first sampling time point (September 2023), number of individuals used in experiments, and the proportion of insects that died or escaped.

| Taxon | Number of insects in mesocosms prior to experiments | Number of insects used in experiments | Proportion (%) of insect mortality or escape |
| --- | --- | --- | --- |
| Water boatmen<br>(*Hesperocorixa* sp.) | 389 | 120 | 70 |
| Backswimmers<br>(*Notonecta* sp.) | 170 | 116 | 32 |
| Diving beetles<br>(*Laccophilus maculosus*) | 215 | 102 | 53 |

To stock the mesocosms, we collected water boatmen, backswimmers and diving beetles in August and September 2023 from two ponds using D-frame nets (Table 1). The Jewkes Pond site is located in Jimtown, Antigonish County, Nova Scotia (45.7072°N, 61.9016°W) and has a surface area of 450-500 m^2^ (Lauff et al., 2025). Griffith Pond site is located in Addington Forks, Antigonish County, Nova Scotia (45.5621°N, 62.1004°W), with an estimated surface area of 350-400 m^2^. Griffith Pond is less than 20 m from the road and thus may have been more saline than Jewkes Pond due to winter salt runoff. Insects collected from field sites were transported back to St. Francis Xavier University (St. FX) campus in coolers filled with water from their respective ponds and transferred to outdoor mesocosms for sampling over the course of our experimental design. We performed additional field collection of insects in late March and early April for direct use in experiments.

### Mesocosm conditions

Mesocosms were located behind the J. Bruce Brown building, on St. FX campus (45.6177°N, 61.9920°W). We used two heavy-duty polyethylene stock tanks to house insects, water and vegetation collected from each respective field site: a Jewkes Pond mesocosm (473 L, 125 gallons) and Griffith Pond mesocosm (379 L, 100 gallons). We also lined the bottom of the stock tanks with aquarium pebbles to provide a substrate that would not influence water pH or increase turbidity. To provide natural insulation and thermal buffering, the stock tanks were placed in the ground at a depth of approximately 0.5 m. To protect insect stocks from terrestrial predators, covers made from PVC piping and two mesh layers (an inner coarse layer and an outer fine layer) were secured over the tanks with zip-ties.

### Outdoor temperature and dissolved oxygen measurements

Natural seasonal variation in water temperature and dissolved oxygen (DO) levels were continuously measured using HOBO U-26-001 oxygen loggers (Hoskins Scientific, Oakville Ontario, Canada) in both Jewkes and Griffith mesocosms, as well as Griffith Pond. Jewkes Pond data is not included due to logger malfunction. To secure loggers in ponds, we zip-tied each logger to a wooden stake that was weighed down by a cement-filled flowerpot. We orientated the loggers so that the sensor end floated just above the substrate, in water at least 30 cm deep. In the mesocosms, the position of the loggers was maintained by tying a rope to a perimeter fence that surrounded the tanks. All loggers were set to record temperature (°C) and DO (mg L^-1^) at 1 h intervals.

### Whole-animal cold tolerance

To measure CT_min_, we used an Arctic A25 recirculating chiller (ThermoFisher) to slowly cool insects in their appropriate mesocosm or pond water until voluntary movement ceased. We processed 12 insects per taxon per time point. We placed each individual in 40 mL of water in a 50 mL polypropylene centrifuge tube, which was held in a custom-made transparent plexiglass chamber with seven other tubes, each also containing an insect (Adams et al., 2025a, Lopez Pedersen et al., 2026). We used an Ecoplus Adjustable Water Pump (EcoPlus, Austin, Texas, USA) to circulate 50% propylene glycol fluid from the chiller into the plexiglass container (Adams et al., 2025a, Lopez Pedersen et al., 2026). The temperature of the propylene glycol fluid was decreased from 4°C at 0.25°C min^-1^ to a temperature at which CT_min_ was reached or the water in the tubes froze for all animals. We measured the temperature of the water in each tube using T-type thermocouples connected to Picolog v6.24.1 software (Pico Technology, Cambridge, UK) via a Pico Technology TC- 08 interface (Toxopeus et al., 2019b). We visually confirmed CT_min_ values when individuals no longer moved voluntarily or responded to physical touch. After this protocol, we briefly dried the insects with a paper towel and weighed them in preparation for measuring SCP.

To measure SCP, insects were transferred into 1.7 mL centrifuge tubes, which were then placed in a custom-made insulated aluminum block that was cooled by a coil containing fluid from the recirculating chiller (Adams et al., 2025b; Toxopeus et al., 2019b). The temperature of the propylene glycol fluid was decreased from 4°C at 0.25°C min^-1^ to a temperature at which all insects froze. Insect body temperature was measured using T-type thermocouples and PicoLog software as described above. The SCP was defined as the lowest temperature observed prior to a large increase in temperature caused by the exothermic process of ice formation (Sinclair et al., 2015). At the termination of SCP experiments, we transferred insects to a −80°C freezer for long-term storage and genetic analysis.

### Whole-animal hypoxia tolerance

We exposed insects to laboratory-simulated hypoxia at each experiment time point to assess behavioural markers of hypoxia tolerance. We used 12 insects from each taxon per time point, separate from those used to measure cold tolerance. Similar to the CT_min_ experimental set up, each insect was kept in 30 mL of mesocosm or pond water in a 50 mL polypropylene tube that was temperature-controlled by circulation of chilled fluid through the custom plexiglass container. Water temperatures were adjusted to mimic that of the mesocosms or ponds during each time point. Animals were kept in these conditions under normoxia for 1 h prior to measuring ASR to acclimate them to the laboratory environment.

To assess ASR, DO in the tube water was decreased to 5% air saturation (1.05 kPa, 0.6 mg L^-1^), and insects had access to normoxic air above the water. P_crit_ values were not available for our study species, so we chose 5% air saturation based on the published P_crit_ for the gill-breathing nymphs of the mayfly, *Neocloeon triangulifer* (Cochran et al., 2022). DO was controlled using a Loligo water parameter controller system (Loligo Systems, Viborg, Denmark). The system consists of four fiber optic oxygen mini sensors connected to a Witrox 4 oxygen meter that relayed the PO_2_ in the water to WitroxCTRL software (Version 2, Loligo Systems) run on a Windows 11 laptop. Once the target DO was reached, Witrox Control software was used to maintain the target DO by controlling relays that opened and closed solenoid switches that regulate air and nitrogen gas delivery to the experimental tubes. Once targeted DO levels were reached and maintained, we recorded the behaviour of two insects in separate tubes simultaneously for 1 h using a GoPro Hero4 camera in a 0.5 s time lapse. We used J Watcher (v1.0) (Blumstein and Daniel, 2007; https://www.jwatcher.ucla.edu/) to process the GoPro footage at 0.5× speed, recording instances of surfacing and leaving the surface, which were then used to calculate the number of surface visits (ASR), time at surface, and time submerged.

### Biochemical assays

To prepare samples for biochemical assays, we weighed and flash-froze 12 individuals per taxon and time point. Insects were stored at −80°C for up to 2 weeks and then homogenized whole in tris-buffered saline (TBS; 5 mM Tris, 137 mM NaCl, KCl 2.7 mM, pH 6.6) using plastic pestles in 1.7 mL tubes on ice (Adams et al., 2025a, Lopez Pedersen et al., 2026). The volume of TBS varied based on general mass differences between insects: 400 μL for backswimmers, 300 μL for water boatmen, and 250 μL for diving beetles. To generate cell-free extracts, we centrifuged homogenates at 3000 × *g* for 5 min at 4 °C, transferred supernatants to new tubes, and centrifuged again at 5000 × *g* for 30 min at the same temperature (Adams et al., 2025a, Lopez Pedersen et al., 2026). Final supernatants were aliquoted into subsets of tubes to be stored at –80°C for each biochemical assay: glucose, glycerol, *myo*-inositol, proline, trehalose, glycogen, and lactate dehydrogenase.

All biochemical assays were conducted using 96-well plates and a SpectraMax iD5 spectrophotometer (Molecular Devices; San Jose, California, USA). All biochemical metrics were standardized to protein concentration, determined using the Pierce BCA Protein Assay Kit (ThermoFisher Scientific) following the manufacturer’s instructions (Pichaud et al., 2025).

Similar to Toxopeus et al. (2019a), we measured whole-insect glucose, glycerol and *myo*-inositol concentrations using the Sigma Glucose (HK) Assay Reagent (Sigma Aldrich, Mississauga, Ontario, Canada), the Glycerol Assay Kit (Megazyme, Bray, Ireland) and the *myo*-Inositol Assay Kit (Megazyme, Bray, Ireland), respectively, according to the manufacturer’s instructions. We measured whole-insect proline and trehalose concentrations following assays described in Carillo and Gibon (2011) and Tennessen et al. (2014), respectively, with modifications described by Toxopeus et al. (2019a). We measured whole-insect glycogen concentrations using methods modified from Mandic et at. (2013). We made two 50 μl aliquots from each sample and split them into two subsets: one subset combined with 10 μL of acetate buffer (sodium acetate, acetic acid) (undigested), and the other with 10 μL of 30 U mL^-1^ amyloglucosidase (Sigma Aldrich) in acetate buffer (digested). These samples were incubated at 37°C for 30 min (Mandic et al., 2013), and then glucose concentration was measured with the Glucose (HK) Assay Reagent (Toxopeus et al., 2019a). The relative amount of glycogen was determined from glucose produced in the amyloglucosidase reaction. We measured whole-insect lactate dehydrogenase (LDH) activity at room temperature (c. 20°C) using a modified and scaled-down assay protocol for microplates, originally based on Bergmeyer and Bernt (1974).

### Data analysis

To determine whether whole-animal and biochemical tolerance to low temperature and hypoxia changed significantly between sampling time points, we conducted statistical analyses in R v4.2.3. We used two-way ANOVAs to determine if all correlates of stress tolerance changed seasonally between sampling intervals and sites (mesocosms) within insect groups, followed by Tukey HSD post-hoc analyses. We used principle component analyses (PCAs) to assess how insect metabolite profiles differed among the sampling times for each insect group.

## Results

### Mesocosm conditions and insect abundance

Mesocosm water temperature and DO concentration were lower in the winter than in the fall. In both Jewkes and Griffith mesocosms, temperature decreased in the late fall, prior to any significant change in DO (Fig. 1). Water temperature stayed below 2°C in both mesocosms for approximately 60 consecutive days after December 28, 2023 (Fig. 1). Similarly, the field site of Griffith Pond stayed below the 2°C threshold for 70 days after January 1, 2024 (Fig. S2). Ice cover was sustained from early January until March (Fig. 1), when air temperatures were consistently below 0°C. Mesocosm water had DO generally above 10 mg L^-1^ (22.7 kPa) during the first three sampling time points (September through January), with some temporary fluctuations below this (Fig. 1). When ice cover was present, DO decreased to approximately 2.0 mg L^-1^ (3 kPa) during March and February in Jewkes and Griffith mesocosms, respectively (Fig. 1). Data logger issues prevented us from determining the minimum DO in Griffith Pond during sustained ice cover (Fig. S2). Insect abundance in mesocosms was sufficiently high to conduct experiments in September, November, and January (except beetles in January). Visual assessment of insects immediately following removal from ice-covered mesocosms in late January confirmed motility. No or few insects remained in the mesocosms in March, so all March measurements were done on field-collected individuals.

**Figure 1.**
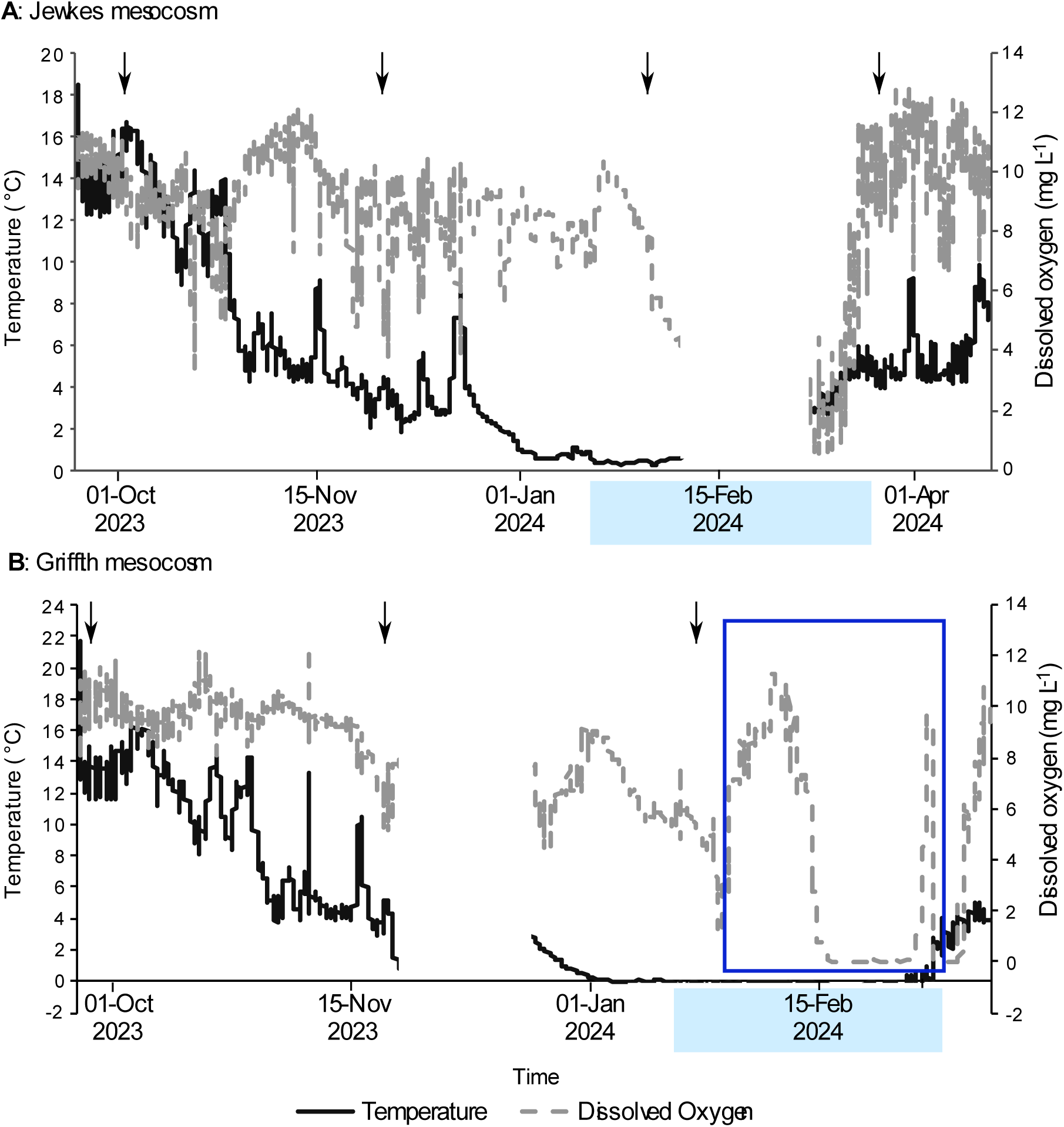
Temperature and dissolved oxygen (DO) concentration recorded in the water of (A) Jewkes and (B) Griffith mesocosms from September 2023 to April 2024. The solid, black line represents temperature, and the grey, dashed line represents DO. Downward-pointing arrows indicate experimental sampling time points. The light-blue rectangles on the x-axes indicate the dates when ice cover was sustained. **(A)** The data gap during February in Jewkes mesocosm occurred due to insufficient logger storage. **(B**) The data gap during November in Griffith mesocosm occurred due to logger transfer between sites. The dark-blue rectangle indicates the period of time during which the logger was encased in ice, and reports conditions in the ice rather than the water.

### Insects remained active and unfrozen under typical pond conditions

When cooled gradually, the temperature at which insects ceased voluntary movement (CT_min_) ranged from 1.3°C to –2.4°C, with the lowest value recorded in water boatmen in November (Fig. 2). For most insect groups, CT_min_ values were only obtained in September 2023, because insects at later time points usually remained active during gradual cooling until the water froze (Fig. 2). The mean freezing temperature of mesocosm water varied over time by as much as 1°C between CT_min_ assays on backswimmers (Fig. 2A; Sampling time *F*_3,35_ = 5.406, *P* = 0.004; Site *F*_1,35_ = 2.360, *P* = 0.133; Sampling time × Site *F*_2,35_ = 1.260, *P* = 0.296) and diving beetles (Fig. 2B; Sampling time *F*_2,18_ = 4.033, *P* = 0.036; Site *F*_1,18_ = 0.008, *P* = 0.932; Sampling time × Site *F*_1,18_ = 0.022, *P* = 0.885), but did not vary among sampling time points for water boatmen (Fig. 2C; Sampling time *F*_3,25_ = 1.609, *P* = 0.212; Site *F*_1,25_ = 0.021, *P* = 0.887; Sampling time × Site *F*_3,25_ = 2.812, *P* = 0.060). When the water froze prior to cessation of movement, the CT_min_ was assumed to be lower than the temperature at which the water froze, but statistical comparison of CT_min_ values over time was therefore not possible for backswimmers and water boatmen. However, several diving beetles had measurable CT_min_ values in November, which were approximately 1°C lower than those measured in September (Fig. 2B; Sampling time *F*_2,22_ = 11.316, *P* < 0.001; Site *F*_1,22_ = 2.151, *P* = 0.156; Sampling time × Site *F*_1,22_ = 0.121, *P* = 0.731).

**Figure 2.**
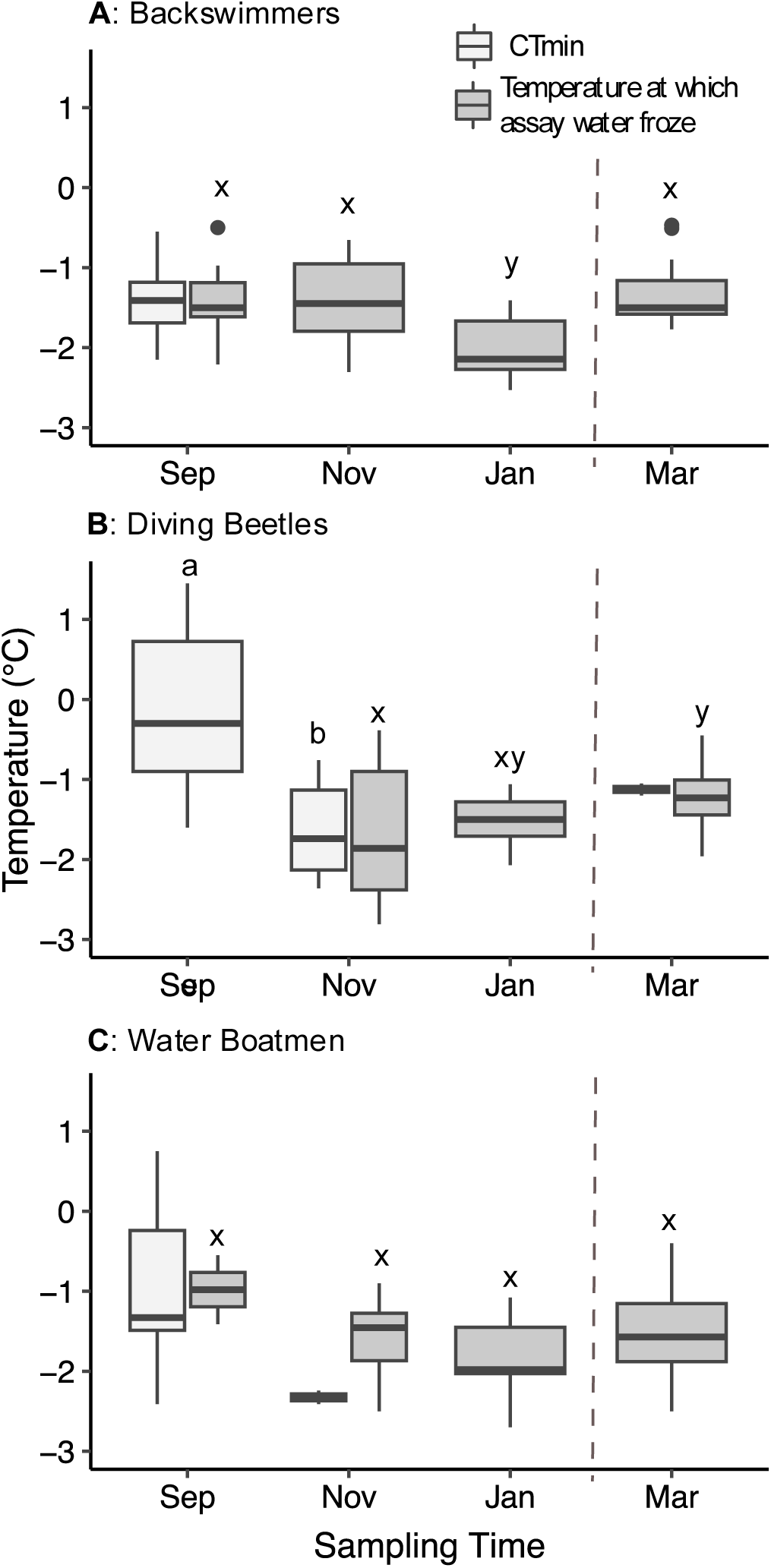
Critical thermal minima (CT_min_; light grey) and assay tube water freezing temperatures (dark grey) of (A) backswimmers (*N* = 50), (B) diving beetles (*N* = 43), and (C) water boatmen (*N* = 52) sampled from Griffith and Jewkes mesocosms in September 2023, November 2023, and January 2024 and from the respective field sites in March 2024. There were 8-12 insects per taxon per time point. No statistical difference was observed between mesocosms, so data was pooled at each time point. The bottom and top of each box represents the lower and upper quartile, respectively; median is represented by the horizontal line; the vertical lines extend to the maximum and minimum values within 1.5 times the inter-quartile range; outliers are indicated by black dots. Different letters above boxes indicate statistical difference between sampling time points, determined using two-way ANOVAs and Tukey’s post-hoc analyses (*P* < 0.05), with a, b used for CT_min_, and x, y used for freezing temperature of assay tube water.

The temperature of spontaneous freezing of internal fluids (SCP; Fig. 3) was consistently lower than the temperature at which water was expected to freeze (Fig. 2). Mean SCP in backswimmers was approximately –7.5°C from September to January and increased by more than 2.5 °C between January mesocosm individuals and March field-collected individuals (Fig. 3A; Sampling time *F*_3,42_ = 4.113, *P* = 0.012; Site *F*_1,42_ = 0.308, *P* = 0.582; Sampling time × Site *F*_3,42_ = 1.491, *P* = 0.231). Mean diving beetle SCP was approximately −10°C from September to January and increased by more than 2.5°C in wild-collected March individuals relative to the previous two sample points (Fig. 3B; Sampling time *F*_3,32_ = 5.641, *P* = 0.003; Site *F*_1,32_ = 0.102, *P* = 0.751; Sampling time × Site *F*_2,32_ = 5.743, *P* = 0.007). Water boatmen mean SCP increased from c. −7°C in September to c. −5°C in November and remained similarly high in January and the field-collected insects in March (Fig. 3C; Sampling time *F*_3,41_ = 10.558, *P* < 0.001, Site *F*_1,41_ = 2.168, *P* = 0.148; Sampling time × Site *F*_3,41_ = 2.764, *P* = 0.054). In pilot studies in fall 2023, we returned frozen insects to chilled pond water for recovery, and none survived (data not shown). Thus, avoiding temperatures below the SCP of these insects is likely important for their overwintering survival.

**Figure 3.**
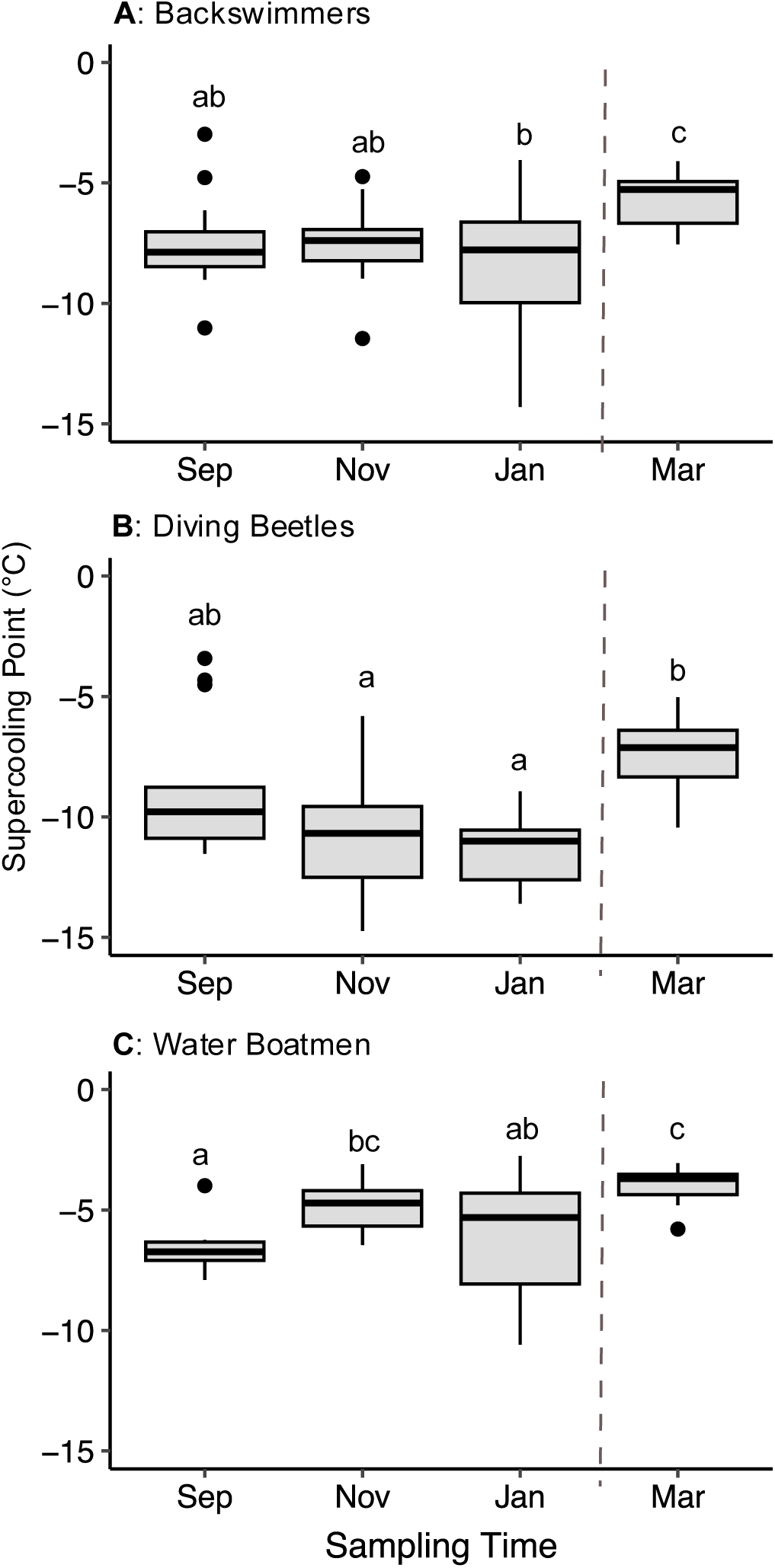
Supercooling point (SCP) temperatures of (A) backswimmers (*N* = 50), (B) diving beetles (*N* = 43), and (C) water boatmen (*N* = 52) sampled from Griffith and Jewkes mesocosms in September 2023, November 2023, and January 2024 and from the respective field sites in March 2024. There were 8-12 insects per taxon per time point. No statistical difference was observed between mesocosms, so data was pooled at each time point. The bottom and top of each box represents the lower and upper quartile, respectively; median is represented by the horizontal line; the vertical lines extend to the maximum and minimum values within 1.5 times the inter-quartile range; outliers are indicated by black dots. Different letters above boxes indicate statistical significance between sampling time points, determined using two-way ANOVAs and Tukey’s post-hoc analyses (*P* < 0.05).

### Whole animal response to hypoxia exhibited minimal seasonal variation

Mean aquatic surface respiration frequency (ASR) did not decrease, and relative time spent submerged did not increase, seasonally during exposures to 1.05 kPa DO (5% air saturation). During low-oxygen exposure, backswimmers approached the water surface relatively infrequently, with typically < 20 visits in 60 min (Fig. 4A), but spent approximately half of their time at the surface across all sampling time points (Table 2; Sampling time *F*_3,44_ = 2.095, *P* = 0.115; Site *F*_1,44_ = 0.048, *P* = 0.828; Sampling time × Site *F*_3,44_ = 0.040, *P* = 0.989). ASR in this group did not change over time, but backswimmers from Jewkes mesocosm visited the water surface approximately 18 more times than those from Griffith mesocosm in January (Fig 4A; Sampling time *F*_3,44_ = 2.107, *P* = 0.113; Site *F*_1,44_ = 0.093, *P* = 0.762 Sampling time × Site *F*_3,44_ = 5.578, *P* = 0.003). Diving beetles spent approximately half of their time at the surface (Table 2; Sampling time *F*_3,37_ = 2.122, *P* = 0.114; Site *F*_1,37_ = 0.479, *P* = 0.493; Sampling time × Site *F*_3,37_ = 0.608, *P* = 0.614), with ASR that ranged from c. 10 to 80 surface visits during the observation period (Fig. 4B), but with no statistical differences in ASR over time or between mesocosms (Sampling time *F*_3,37_ = 0.412, *P* = 0.745; Site *F*_1,37_ = 0.096, *P* = 0.758; Sampling time × Site *F*_3,37_ = 2.336, *P* = 0.089). Conversely, water boatmen spent most (>75%) of their time submerged across all sampling times (Table 2; Sampling time *F*_3,39_ = 0.571, *P* = 0.637; Site *F*_1,39_ = 1.600, *P* = 0.213; Sampling time × Site *F*_3,39_ = 0.407, *P* = 0.749), with some individuals exhibiting the highest ASRs in this study at more than 80 surface visits in 60 min (Fig. 4C). ASR in water boatmen differed over time and between mesocosms, with the highest values observed in Griffith mesocosm insects during November (Fig. 4C; Sampling time *F*_3,40_ = 9.555, *P* < 0.001; Site *F*_1,40_ = 20.426, *P* < 0.001; Sampling time × Site *F*_3,40_ = 4.126, *P* = 0.012).

**Figure 4.**
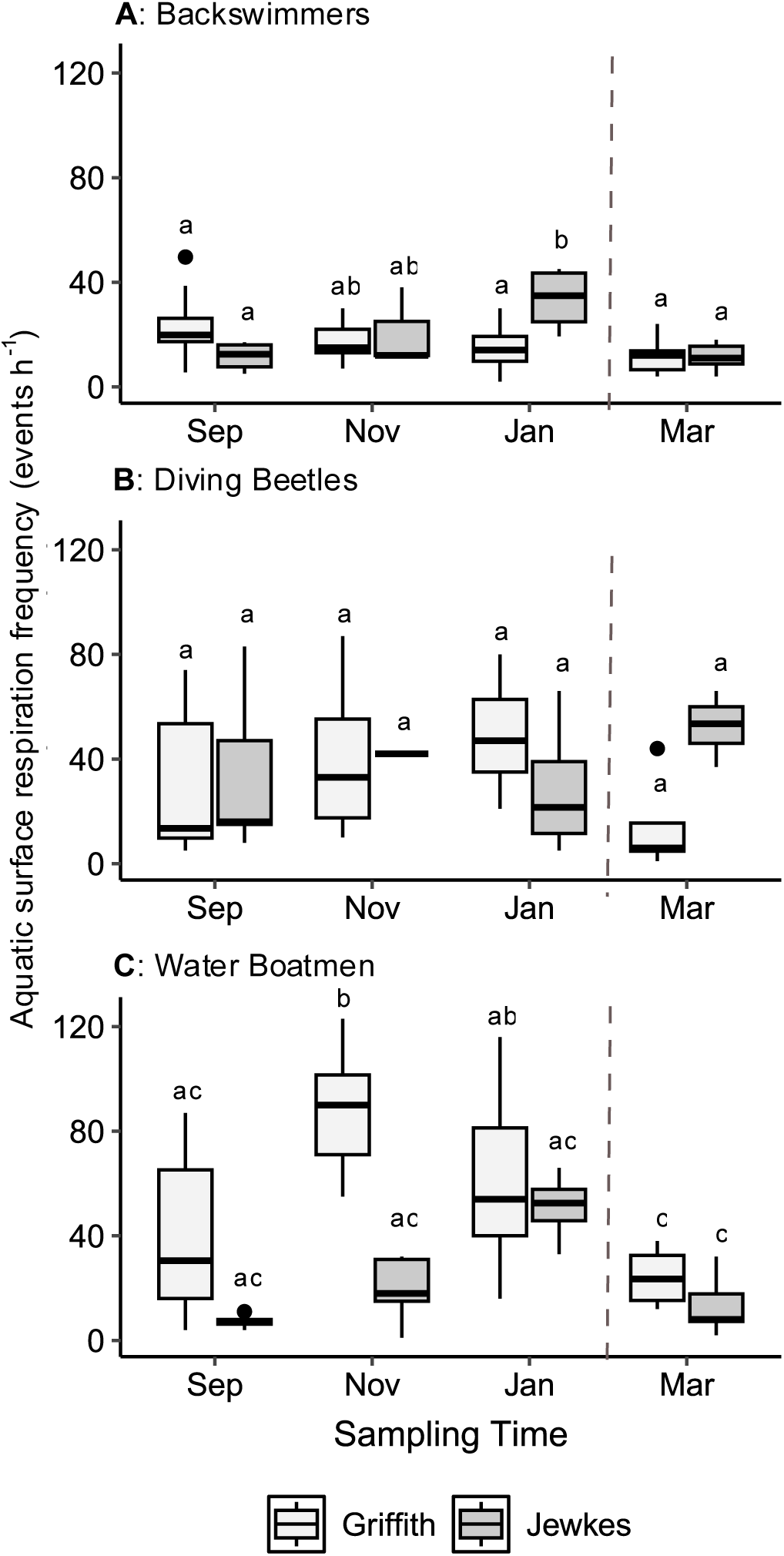
Frequency of surface visits for respiration during 1 h exposure to 5% air saturation (1.05 kPa, 0.6 mg L^-1^) by (A) backswimmers (*N* = 52), (B) diving beetles (*N* = 45), and (C) water boatmen (*N* = 48) sampled from Griffith (light grey) and Jewkes (dark grey) mesocosms in September 2023, November 2023, and January 2024 and from the respective field sites in March 2024. There were 4-8 insects per taxon per mesocosm at each time point. The bottom and top of each box represents the lower and upper quartile, respectively; the median is represented by the horizontal line; the vertical lines extend to the maximum and minimum values with 1.5 times the inter-quartile range; outliers are indicated by black dots. Different letters above boxes indicate statistical significance between sampling time points or mesocosm, determined using two-way ANOVAs and Tukey’s post-hoc analyses (*P* < 0.05)

**Table 2.** Mean (*± SE)* time in minutes spent at water surface during a 60 min exposure to 1.05 kPa oxygen (5% air saturation) by backswimmers, diving beetles, and water boatmen sampled from mesocosms in September 2023, November 2023, and January 2024 and from the field in March 2024. There was no significant difference in time spent at the water surface over time or between mesocosms within each taxon (two-way ANOVAs, see text).

| Sampling time | Backswimmers | Diving beetles | Water boatmen |
| --- | --- | --- | --- |
| September | 31.0 ± 4.8 (n = 16) | 40.3 ± 3.9 (n = 16) | 5.2 ± 2.2 (n = 12) |
| November | 25.1 ± 3.9 (n = 12) | 29.4 ± 5.3 (n = 9) | 8.6 ± 1.9 (n = 11) |
| January | 34.7 ± 2.6 (n = 12) | 31.5 ± 3.7 (n = 12) | 9.6 ± 2.1 (n = 12) |
| March<br>(field-collected) | 19.9 ± 4.5 (n = 12) | 23.7 ± 7.3 (n = 8) | 9.3 ± 3.8 (n = 12) |

### Biochemical correlates of stress tolerance had species-specific patterns

Each species exhibited different seasonal patterns in the concentrations of low molecular weight metabolites in whole-body homogenates. In backswimmers, mean total concentrations of potential cryoprotectants varied seasonally and between sites (Fig. 5A; Sampling time *F*_3,42_ = 52.628, *P* < 0.001; Site *F*_1,42_ = 4.411, *P* = 0.042; Sampling time × Site *F*_3,42_ = 5.567, *P* = 0.003).

**Figure 5.**
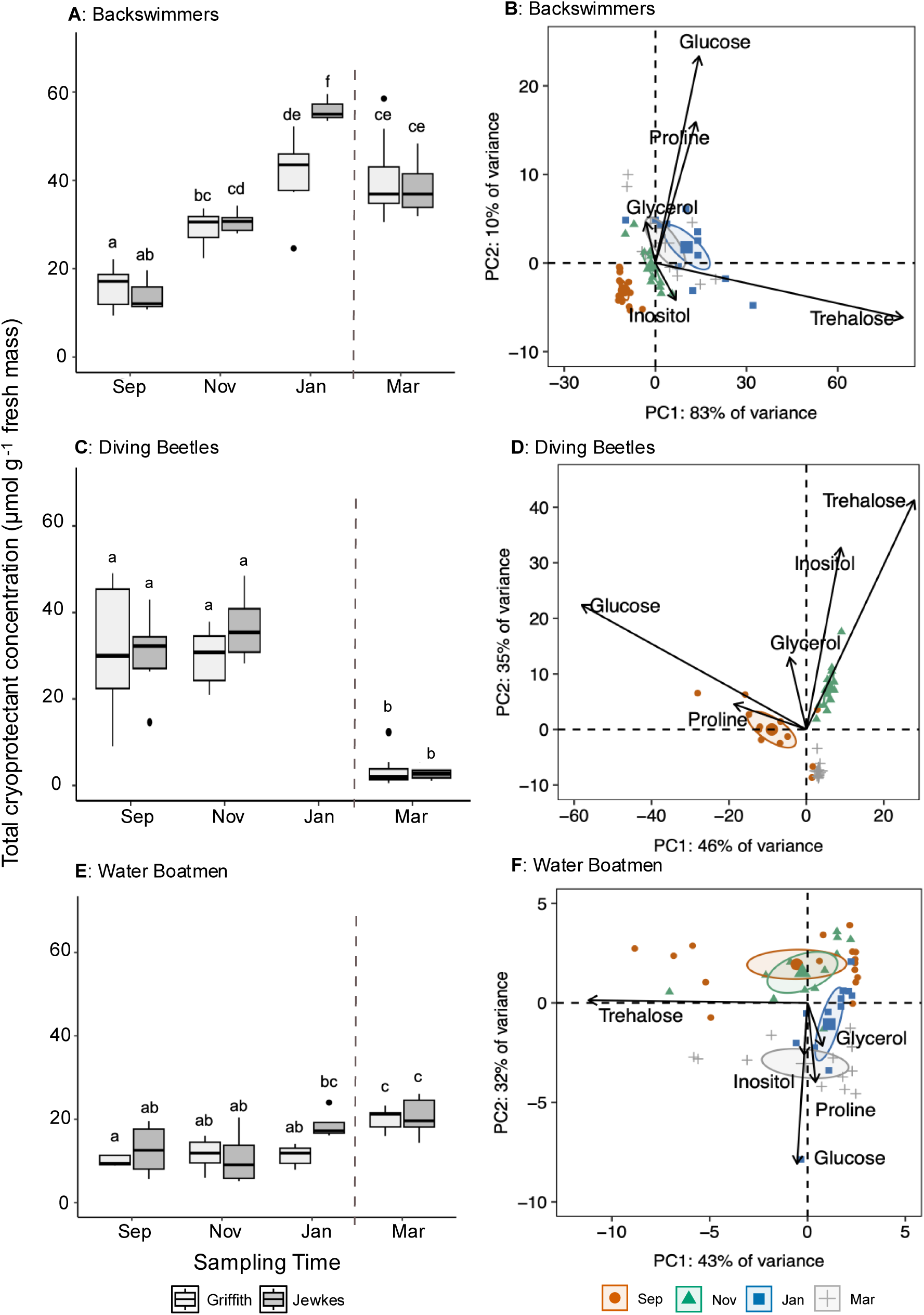
Low molecular weight metabolites profiles determined from whole body homogenates of (A, B) backswimmers (*N* = 50), (C, D) diving beetles (*N* = 36), and (E, F) water boatmen (*N* = 52) sampled from Griffith and Jewkes mesocosms in September 2023, November 2023, and January 2024 and from the respective field sites in March 2024. There were 4-8 insects per taxon per mesocosm at each time point. **(A, C, E)** Sum of the concentrations of all five low molecular weight metabolites (glycerol, glucose, *myo*-inositol, proline, and trehalose) measured via spectrophotometric assays. Different letters above boxes indicate statistical significance between sampling time points or mesocosm, determined using two-way ANOVAs and Tukey’s post-hoc analyses (*P* < 0.05). **(B, D, F)** Principle components analyses (PCAs) based on the concentrations of the same five low molecular weight metabolites. Each small point represents one insect, while means of each group are represented by the larger points. The circles represent a 95% confidence interval. Arrows represent the influence of metabolite on position points within the coordinate space; the length of the arrow corresponds with the magnitude of influence. Concentrations of individual cryoprotectants are available in Figs. S3-5.

These concentrations were about two-fold higher in November and more than three-fold higher in January compared to September but did not differ between January sampling and the wild-caught March backswimmers (Fig. 5A). Differences were driven largely by high concentrations of glucose and proline in January and March (Fig. 5B, Fig S3). Mean total metabolite concentration was higher in backswimmers from Jewkes than Griffith mesocosm in January, but statistically the same between mesocosms at other time points (Fig. 5A). In diving beetles, total metabolite concentrations were approximately 30 μmol g^-1^ in both September and November but were about six-fold lower in wild-caught March beetles, and did not differ between sites (Fig. 5C; Sampling time *F*_2,30_ = 39.588, *P* < 0.001; Site *F*_1,30_ = 0.392, *P* = 0.536; Sampling time × Site *F*_2,30_ = 0.799, *P* = 0.459). All five metabolites influenced differences over time in diving beetles (Fig. 5D), with a shift from high glucose, glycerol, and proline concentrations in September to high glycerol, *myo*-inositol, and trehalose concentrations in November, and low concentrations of all metabolites from field-collected beetles in March (Fig. S4). The total concentration of metabolites in water boatmen was highest in wild-caught insects in March, and there were minor differences between insects from Jewkes and Griffith mesocosms (Fig. 5E; Sampling time *F*_3,44_ = 13.956, *P* < 0.001; Site *F*_1,448_ = 4.612, *P* = 0.037; Sampling time × Site *F*_3,44_ = 2.216, *P* = 0.010). Trehalose concentrations varied substantially among water boatmen within each sampling group, while *myo*-inositol, proline, and glucose all exhibited increases in concentration as the season progressed (Fig. 5F, Fig. S5).

Glycogen concentrations changed seasonally in backswimmers and diving beetles, but not in water boatmen (Fig. 6). In backswimmers, mean glycogen concentrations decreased approximately two-fold between September and January; and glycogen concentrations were similar in field-collected March backswimmers and mesocosm individuals in January (Fig. 6A; Sampling time *F*_3,42_ = 4.386, *P* = 0.009; Site *F*_1,42_ = 1.549, *P* = 0.220; Sampling time × Site *F*_3,42_ = 1.399, *P* = 0.256). In diving beetles, glycogen concentrations were statistically similar between both sites at each time point, and but were about seven-fold lower in all wild insects in March relative to November mesocosm beetles (Fig. 6B; Sampling time *F*_2,30_ = 5.468, *P* = 0.009; Site *F*_1,30_ = 0.215, *P* = 0.646; Sampling time × Site *F*_2,30_ = 0.157, *P* = 0.855). Glycogen concentrations did not change seasonally or between mesocosms in water boatmen and was highly variable among individuals (Fig. 6C; Sampling time *F*_3,44_ = 1.657, *P* = 0.190; Site *F*_1,44_ = 0.009, *P* = 0.932; Sampling time × Site *F*_3,44_ = 3.370, *P* = 0.027).

**Figure 6.**
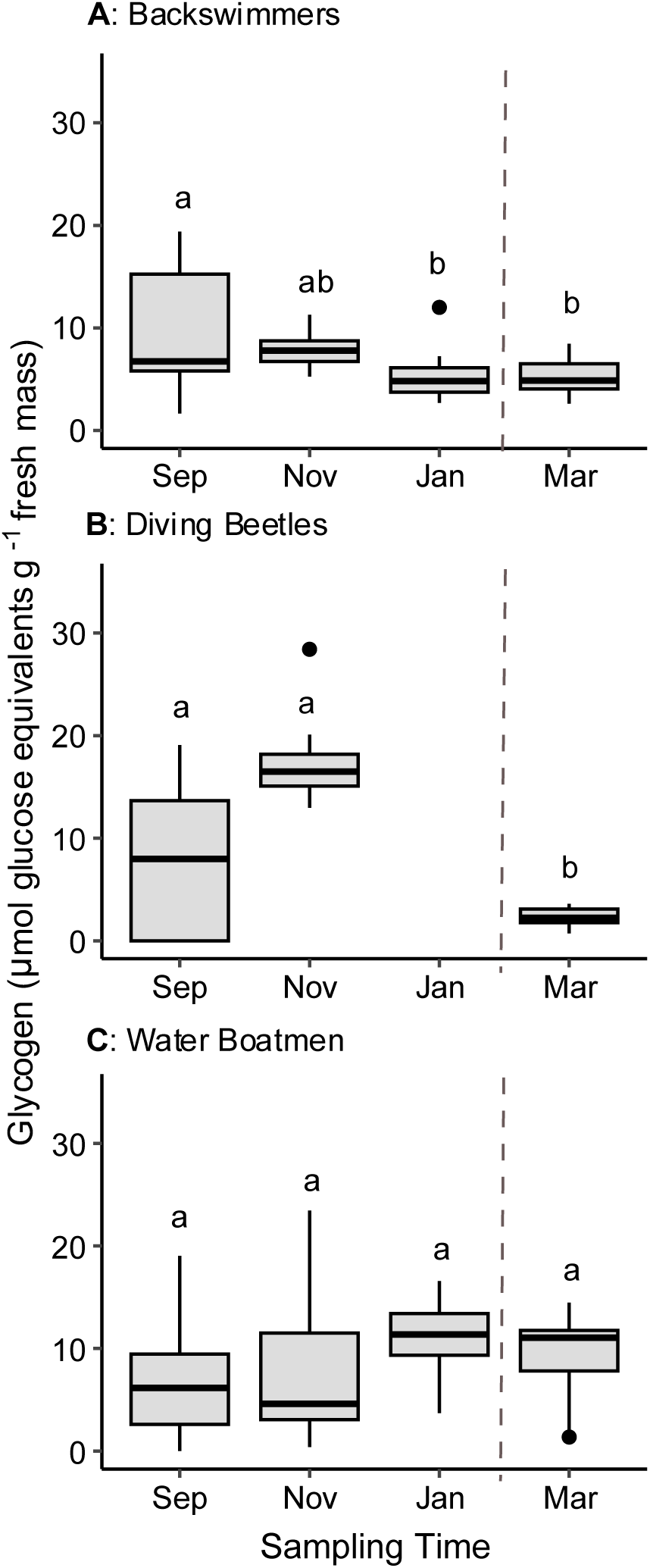
Spectrophotometrically-determined relative concentrations of glycogen from whole body homogenates of (A) backswimmers (*N* = 50), (B) diving beetles (*N* = 36), and (C) water boatmen (*N* = 52) sampled from Griffith (light grey) and Jewkes (dark grey) mesocosms in September 2023, November 2023, and January 2024 and from the respective field sites in March 2024. There were 8-12 insects per taxon per time point. No statistical difference was observed between mesocosms, so data was pooled at each time point. The bottom and top of each box represents the lower and upper quartile, respectively; the median is represented by the horizontal line; the vertical lines extend to the maximum and minimum values with 1.5 times the inter-quartile range; outliers are indicated by black dots. Different letters above boxes indicate statistical significance between sampling time points, determined using two-way ANOVAs and Tukey’s post-hoc analyses (*P* < 0.05).

Whole animal lactate dehydrogenase (LDH) activity exhibited small seasonal changes in backswimmers and water boatmen, but not in diving beetles (Fig. 7). In backswimmers, LDH activity decreased mildly between September and November and January, and did not differ between sites (Fig. 7A; Sampling time *F*_3,42_= 4.225, *P* = 0.011; Site *F*_1,42_= 3.095, *P* = 0.086; Sampling time × Site *F*_3,42_ = 1.166, *P* = 0.334). There were no statistical differences between wild-caught March insects and captive mesocosm insects from the previous three time points (Fig. 7A). LDH activity in diving beetles did not change between sampling points or mesocosms and was in a similar range to backswimmers (Fig. 7B; Sampling time: *F*_2,30_ = 1.366, *P* = 0.270; Site *F*_1,30_ = 0.025, *P* = 0.876; Sampling time × Site *F*_2,30_ = 0.654, *P* = 0.527). LDH activity was about 10-fold lower in water boatmen compared to the other two insect taxa, and did not differ between mesocosms or from September to January (mean 0.14 μmol min^-1^ μg^-1^ protein), but was about two-fold higher in wild-caught individuals in March (0.34 μmol min^-1^ μg^-1^ protein) (Fig. 7C; Sampling time *F*_3,44_ = 4.668, *P* = 0.006; Site *F*_1,44_ = 3.415, *P* = 0.071; Sampling time × Site *F*_3,44_ = 1.044, *P* = 0.383).

**Figure 7.**
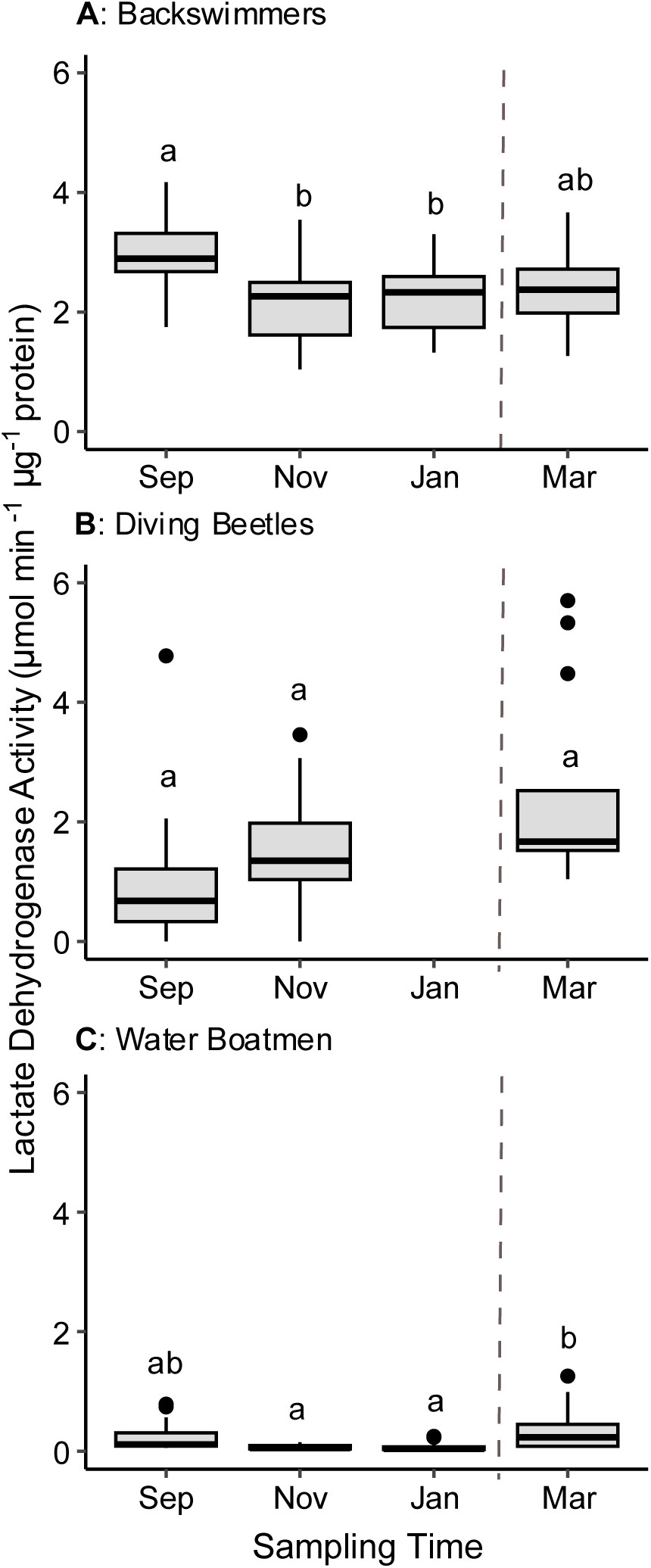
Spectrophotometrically-determined lactate dehydrogenase activity measured in whole body homogenates from (A) backswimmers (*N* = 50), (B) diving beetles (*N* = 36), and (C) water boatmen (*N* = 52) sampled from Griffith and Jewkes mesocosms in September 2023, November 2023, and January 2024 and from the respective field sites in March 2024. There were 8-12 insects per taxon per time point. No statistical difference was observed between mesocosms, so data was pooled at each time point. The bottom and top of each box represents the lower and upper quartile, respectively; median is represented by the horizontal line; the vertical lines extend to the maximum and minimum values with 1.5 times the inter-quartile range; outliers are indicated by black dots. Different letters above boxes indicate statistical significance between sampling time points, determined using two-way ANOVAs and Tukey’s post-hoc analyses (*P* < 0.05).

## Discussion

This research has expanded our understanding of the seasonal stress physiology of aquatic insects overwintering in temperate ponds, as well as the environmental conditions that occur before and during sustained ice cover. We obtained partial support for our hypothesis that cold tolerance would increase seasonally. Water temperature of both our mesocosms and ponds decreased to near 0°C as air temperature dropped in the fall, which was associated with a decrease in diving beetle CT_min_ values. The SCPs of all three taxa – backswimmers, diving beetles, and water boatmen – were several degrees below the temperature of the water, allowing insects to remain unfrozen and motile throughout the winter. Only one taxon (backswimmers) showed an overall increase in the concentration of low molecular weight metabolites as the season progressed, which occurred with depletion of glycogen stores. However, the concentrations of several putative cryoprotectants (glucose, glycerol, *myo*-inositol, proline, and trehalose) were fairly moderate across all three taxa, and the extent to which these molecules affect cold tolerance is unclear. We obtained no support for our hypothesis that hypoxia tolerance would improve to facilitate survival in ice-covered aquatic environments. The mesocosm water had mild oxygen fluctuations from September through January, and the insects sampled from these mesocosms did not decrease ASR or increase their ability to stay submerged when exposed to hypoxia in the laboratory, nor did they increase LDH activity. Glycogen reserves declined seasonally only in backswimmers, but we suggest that this depletion largely drove metabolite synthesis, rather than supporting catabolism for anaerobic ATP production (Hochachka et al., 2022; Clark and Worland, 2008). Regrettably, after mesocosms DO decreased substantially (3 kPa, 2 mg L^-1^) later in the season there were no more mesocosm insects to sample, so we could not determine the effect of the hypoxic mesocosm environment on our study species.

### Winter insects remained active until water froze and did not depress SCP

Although we could not directly measure CT_min_ in most of our overwintering insects, we infer that it was generally below 0°C because most insects remained active until water froze in November through March. The values we measured were similar to those of high-altitude Colorado stonefly, mayfly and caddisfly larvae (near or below the freezing temperature of water; Shah et al., 2017), and lower than those measured in damselfly nymphs in the United Kingdom (c. 2 to 9°C; Smith and Lancaster, 2020). One study has recorded lower CT_min_ values (c. −4°C) for water boatmen and diving beetles in Chile (Rendoll-Cárcamo et al., 2020), suggesting their CT_min_ protocol allowed the environmental water to stay liquid to lower temperatures than in our study. We speculate that the true CT_min_ of the winter insects in our study is below –2°C, based on comparisons to the limited studies that have successfully measured CT_min_ in water at subzero temperatures. Future work could examine the mechanisms underlying successful motility in near-freezing water, such as regulation of ion gradients necessary for neuromuscular coordination (e.g., Andersen et al., 2015). The ability to remain motile during the winter may allow these insects to continue foraging and maintain the energy reserves needed for basic bodily functions and stress tolerance mechanisms.

Our results suggest that depression of SCP via cryoprotectant accumulation is not important for overwintering survival of our aquatic insects. SCP did decrease prior to or during winter, despite moderate increases in the concentrations of putative cryoprotectants (e.g., in the backswimmers). SCPs ranged from c. −4°C to –11°C, which was similar to other aquatic insects (Carbonell et al., 2024), but higher than freeze-avoidant terrestrial insects (Khajehoseini Saleh Abad et al., 2023; Lopez Pedersen et al., 2026; Watanabe et al., 2002), which experience much lower temperatures during winter than aquatic insects. Concentrations of putative cryoprotectants were also low compared to terrestrial insects that depress their SCP (Khani et al., 2007; Saeidi and Moharramipour, 2017). Depressing SCP in the winter may be unnecessary for aquatic insects due to the relatively high temperature of unfrozen water, but proximity to ice still poses risks. Water boatmen in Saskatchewan, Canada have survived encasement in pond ice, although do not appear to be freeze-tolerant (Srayko et al., 2023). Future work could examine whether aquatic insects accumulate antifreeze molecules (Box et al., 2022; Duman, 2015; Guz et al., 2014; Hawes et al., 2011; Tyshenko et al., 1997) to avoid freezing when in contact with, or encased by, ice.

### Seasonal changes in hypoxia stress tolerance were challenging to detect

The relatively mild behavioural response observed under hypoxia in this study is consistent with previous studies demonstrating that air-breathing insects can tolerate prolonged exposure to severe hypoxia. For example, air-breathing insects (e.g., *Limnius volckmari, Aphelocheirus aestivalis*) are generally less vulnerable to high temperature stress under hypoxia compared to gill breathers (e.g., *Rhitrogena semicolorata*, *Calopteryx virgo*), maintaining a higher critical thermal maximum, CT_max_ (Verberk and Bilton, 2013). At 10°C, diving beetles (*Platynectes decempunctatus*) can approximately double their dive durations compared to those at 20°C, partly due to lower metabolic (oxygen) demand (Jones and Seymour, 2021). Thus, the low temperatures of the water used in our experiments may have facilitated a low dependence on atmospheric oxygen. Future work could compare our observations to summer-acclimatized insects to fully evaluate whether the diving behaviour observed here represents a winter-specific response or is similar to the summer baseline patterns for our study species. However, the limited behavioural responses observed in the present study may also reflect the relatively mild hypoxic challenge imposed by our experimental design.

We evaluated diving and surfacing behaviour under conditions where the water was hypoxic (1.1 kPa PO_2_), but the atmosphere remained normoxic, allowing insects unrestricted access to atmospheric oxygen throughout the experiment. As a result, the laboratory assay only challenged the aquatic component of their bimodal respiratory strategy (Seymour and Matthews, 2013), while aerial respiration remained unaffected. This differs from the conditions in ice-covered ponds, where ice and snow can restrict or completely block access to the air-water interface (Jansen et al., 2025; Mathias and Barica, 1980; Zhang et al., 2024). Furthermore, as the insects had access to normoxic air throughout the experiment, oxygen depletion within their physical gill during the 1 h exposure may not have been sufficient to substantially impair gas exchange or elicit measurable behavioural ASR responses. For example, tiger beetle larvae (*Cicindela* spp.) can survive for as long as 9 to 22 h when exposed to severe hypoxia (PO_2_ 0.68 kPa, 20°C; Brust and Hoback, 2009), substantially longer than the 1 h exposure used in the present study. Given the limited physiological data available for aquatic insects (Cochran et al., 2022), future studies should determine baseline hypoxia tolerance thresholds such as critical oxygen tension (P_crit_) and incipient lethal oxygen saturation (ILOS) under controlled laboratory conditions. These baseline measurements would provide a foundation for examining how hypoxia tolerance changes under more ecologically relevant scenarios, including prolonged hypoxia and restricted access to atmospheric oxygen.

In contrast to our predictions, there was very little biochemical evidence (glycogen depletion, LDH activity increase) that the aquatic insects used in this study were reliant on anaerobic metabolic in response to declines in environmental oxygen in the mesocosms. One explanation is that, despite declining environmental oxygen, the insects did not experience functional hypoxia (Harrison et al., 2018). For example, a relatively hypoxia-sensitive mayfly larvae (*Neocloeon triangulifer*) exhibited no changes in transcript abundance of LDH and EGL-9 (prolyl hydroxylase) at low dissolved oxygen concentrations that were above the P_crit_ value for this species, but upregulated both hypoxia-responsive genes below the P_crit_ (Cochran et al. 2022). Several physiological factors may facilitate suppression of the P_crit,_ including the use of haemoglobin (present in at least one backswimmer species, *Anisops deanei*; Matthews and Seymour, 2008; Miller, 1964; Wawroski et al., 2012) and remodelling of the tracheal system to improve oxygen delivery to tissues (reviewed by Centanin et al., 2010). Thus, declining environmental oxygen does not necessarily result in functional hypoxia, and it is important to determine how behavioural, respiratory, and physiological mechanisms may support tissue oxygenation and maintain aerobic metabolism in these environments to better understand overwintering of aquatic insects.

### Potential mechanisms of metabolite accumulation and their functions

Unlike many terrestrial overwintering insects, our three aquatic taxa did not accumulate large concentrations of putative cryoprotectants to decrease their SCP, and we therefore speculate there are other functions for the low molecular weight metabolites we quantified in our study. First, many of the metabolites we measured (trehalose, glucose, proline, glycerol) can be used as metabolic fuels to generate ATP (Jørgensen et al., 2021; Pichaud et al., 2025; Teulier et al., 2016; Wan et al., 2014) in the potentially energy-limited winter pond environment. For example, diving beetles may consume proline to fuel muscle function and motility (similar to other coleopterans; Jørgensen et al., 2021; Teulier et al., 2016), resulting in the decrease in proline concentration in the fall. Similarly, glycerol concentration decreased in water boatmen during fall and winter, indicating it may have also been consumed as metabolic fuel or used in the synthesis of lipids (Chew and Than, 2021; Lopez Pedersen et al., 2026; Van der Horst et al., 1983). Secondly, cryoprotectants such as glycerol, *myo*-inositol, trehalose and proline can have stabilizing effects on proteins and cell membranes, even at relatively moderate concentrations (Grgac et al., 2022; Toxopeus et al., 2019a; Watanabe, 2002). Thus, the seasonal accumulation of proline in water boatmen and backswimmers, *myo*-inositol in diving beetles and water boatmen, and trehalose in backswimmers could perform some cryoprotective functions.

Unlike many terrestrial insects that overwinter in diapause and do not feed (Sinclair, 2015; Toxopeus et al., 2024), the insects in our study remained motile during winter and could potentially accumulate metabolites through their diet. For example, dietary proline accumulation has been described in drosophilid species during cold acclimation (Moos et al., 2022). Backswimmers may not have had access to their normal prey under winter ice, as they are primarily surface predators (Sano and Kurokura, 2011), so their increase in total metabolite concentrations may have been derived from stored glycogen, which was depleted by about 50% from September and January. As more generalist scavengers and detritovores (Polhemus, 2008; White and Roughley, 2008), the diving beetles and water boatmen likely had relatively more reliable access to food than backswimmers in the mesocosms during the winter and thus could accumulate various metabolites without depleting glycogen. However, the very low concentration of glycogen and total metabolites in wild-caught diving beetles in March suggest that the natural pond environment may be considerably energy-limited. Future work that examines the effect of ecological niche and diet on metabolite accumulation in aquatic insects during the winter could expand our understanding of this system.

### Mesocosms as a model for simulating temperate ponds

The use of mesocosms rather than seasonal field collections presented a number of benefits and limitations that should be considered when planning future work. Importantly, temperature and DO in mesocosms were often comparable with measurements taken from the field, although we lack data for the Griffith Pond for the most hypoxic time (February) in the Griffth mesocosm due to issues with our data logger. However, it is uncertain if the overall stress being experienced by insects in these environments differed, as pond sites were larger and more complex than the mesocosms, with the presence of water inflow and outflow, and potential differences in snow and ice quality, humic content (dissolved organic carbon), microbial activity, seasonal carryover effects of summer productivity, and differences in natural buffering capacity (Danks, 2008; Gorsky et al., 2024; Jansen et al., 2025; Mathias and Barica, 1980; Srayko et al., 2023). The greatest benefit of mesocosm use was the convenience of sampling from an abundant, pre-collected pool of insects on campus during winter, when it is challenging and labour-intensive to sample insects directly from ice-covered ponds. However, we still saw substantial insect mortality or escape in our mesocosm system, and it is unclear how much of this was normal (i.e., would occur in ponds) or artificially induced by mesocosm conditions. For example, the mortality among water boatmen could be attributed to several factors, including predation by backswimmers, insufficient or low-quality nutrition, competition, life cycle, and general stress resulting from movement to a new and unnatural habitat (Olkeba et al., 2021).

### Conclusions

Our study shows that multiple air-breathing insect taxa exhibit adaptations that facilitate overwintering in ice-covered ponds, expanding our understanding of the stress physiology of organisms in these important but understudied habitats. We demonstrated that backswimmers, diving beetles, and water boatmen were able to stay motile at low temperatures in the winter and exhibited different seasonal patterns of SCP and cryoprotectant concentrations than terrestrial freeze-avoidant insects. We characterized dive behaviour under acute hypoxia exposure, as well as seasonal patterns of biochemical metrics associated with hypoxia tolerance, in our three taxa of air-breathing aquatic insects, but more research is needed to fully understand hypoxia stress and stress tolerance in ice-covered ponds. Future research should investigate cold and hypoxia stress both separately and together to better understand how stress responses interact with each other in an ecologically relevant context.

## Acknowledgements

The authors would like to thank the StFX Animal Care Facility and Facilities Management for assistance setting up mesocosms; T.M. Clow, A.L. Gough, J. Lopez Pedersen, S.E. Rokosh, and M.L. van Oirschot for assistance with field work; B.R. Taylor and R.F. Lauff for assistance with insect identification; R.C. Wyeth and T. Llewellyn for assistance with processing behavioural video data, and the landowners who provided permission to collect insects from their ponds: the Griffith family, L. Jewkes, B. MacKinnon, and the Sisters of St. Martha congregation.

## Contributions

**LSB**: conceptualization, data curation (non-genetic), formal analysis, funding acquisition, investigation (non-genetic), methodology, writing – original draft, writing – review & editing. **TAK**: data curation (genetic), funding acquisition, investigation (genetic), methodology, writing – original draft, writing – review & editing. **TMR**: conceptualization, supervision, writing – original draft, writing – review & editing. **JT**: conceptualization, funding acquisition, supervision, writing – original draft, writing – review & editing.

## Data availability

All data and code used to analyse the data are available from Zenodo: https://doi.org/10.5281/zenodo.21480292. Genetic sequences are deposited in NCBI (https://www.ncbi.nlm.nih.gov/), accession numbers PZ723897 - PZ723906 (Table S1).

