## Supplementary Material for "Overwintering under the ice: seasonal shifts in cold and hypoxia stress physiology of three winter-active pond insects"

*corresponding author

**Table S1**. Likely identity of water boatmen (Hesp), backswimmer (Noto), and diving beetle (Lacc) samples collected from two ponds in Eastern Nova Scotia, based on genetic similarity of partial *COI* (*cytochrome oxidase subunit I*) to sequences in NCBI (National Centre for Biotechnology Information) and species currently documented in Nova Scotia.

| Sample from this study | | Top BLAST Hit in NCBI | | | Nova Scotian[a] species within BLAST Hit genus | Likely identity of samples |
|---|---|---|---|---|---|---|
| ID | NCBI Accession Number | Species (location[a]) | Accession number | Genetic Identity | | |
| Hesp_01 | PZ723897 | *Hesperocorixa lucida* (Ontario, Canada; Illinois and Ohio, USA) | OP849950.1 | 91.9 % | *H. interrupta*, *H. minorella*, *H. kennicotti* | *Hesperocorixa* sp. |
| Hesp_02 | PZ723898 | | OP849950.1 | 91.9 % | | |
| Hesp_04 | PZ723899 | | OP849950.1 | 91.9 % | | |
| Noto_05 | PZ723900 | *Notonecta amplifica* (Asia) | MZ305077.1 | 89.4 % | *N. borealis*, *N. insulata*, *N. irrorata*, *N. lunata*, *N. undulata* | *Notonecta* sp. |
| Noto_06 | PZ723901 | | MZ305077.1 | 89.4 % | | |
| Noto_07 | PZ723902 | | MZ305077.1 | 89.4 % | | |
| Noto_08 | PZ723903 | | MZ305077.1 | 89.4 % | | |
| Lacc_09 | PZ723904 | *Laccophilus maculosus* (Eastern and Western Canada and USA) | DQ112647.1 | 99.8% | *L. maculosus* | *L. maculosus* |
| Lacc_10 | PZ723905 | | DQ112647.1 | 99.8% | | |
| Lacc_11 | PZ723906 | | DQ112647.1 | 99.8% | | |

[a]Species location information determined from BOLD (Barcode of Life Database; https://www.boldsystems.org/) and GBIF (Global Biodiversity Information Facility; https://www.gbif.org/)

Figure S1 page 1/3

**A: *Hesperocorixa* sp.**

```
Hesp_01    GAACAAAAATTAATTACTCACCATCAATTTTATGGGCCCTGGGATTTGTATTCCTATTTA  60
Hesp_04    GAACAAAAATTAATTACTCACCATCAATTTTATGGGCCCTGGGATTTGTATTCCTATTTA  60
Hesp_02    GAACAAAAATTAATTACTCACCATCAATTTTATGGGCCCTGGGATTTGTATTCCTATTTA  60
           ************************************************************

Hesp_01    CAGTGGGAGGATTAACTGGAGTAGTCCTAGCAAACTCGTCAATTGATATTGTAATACACG  120
Hesp_04    CAGTGGGAGGATTAACTGGAGTAGTCCTAGCAAACTCGTCAATTGATATTGTAATACACG  120
Hesp_02    CAGTGGGAGGATTAACTGGAGTAGTCCTAGCAAACTCGTCAATTGATATTGTAATACACG  120
           ************************************************************

Hesp_01    ATACATATTATGTAGTTGCACATTTCCACTATGTACTTTCAATAGGAGCTGTATTTGCTA  180
Hesp_04    ATACATATTATGTAGTTGCACATTTCCACTATGTACTTTCAATAGGAGCTGTATTTGCTA  180
Hesp_02    ATACATATTATGTAGTTGCACATTTCCACTATGTACTTTCAATAGGAGCTGTATTTGCTA  180
           ************************************************************

Hesp_01    TTATTGGTAGATTTATTCAATGATACCCATTATTTACAGGACTATCATTAAATCCAAAAT  240
Hesp_04    TTATTGGTAGATTTATTCAATGATACCCATTATTTACAGGACTATCATTAAATCCAAAAT  240
Hesp_02    TTATTGGTAGATTTATTCAATGATACCCATTATTTACAGGACTATCATTAAATCCAAAAT  240
           ************************************************************

Hesp_01    GATTAAAGATTCACTTCATAATTATATTCGTAGGAGTAAATACAACATTTTTCCCTCAGC  300
Hesp_04    GATTAAAGATTCACTTCATAATTATATTCGTAGGAGTAAATACAACATTTTTCCCTCAGC  300
Hesp_02    GATTAAAGATTCACTTCATAATTATATTCGTAGGAGTAAATACAACATTTTTCCCTCAGC  300
           ************************************************************

Hesp_01    ATTTTTTAGGATTAAGAGGAATACCTCGACGGTATTCAGACTACCCTGATAATTTTACAA  360
Hesp_04    ATTTTTTAGGATTAAGAGGAATACCTCGACGGTATTCAGACTACCCTGATAATTTTACAA  360
Hesp_02    ATTTTTTAGGATTAAGAGGAATGCCTCGACGGTATTCAGACTACCCTGATAATTTTACAA  360
           ********************** *************************************

Hesp_01    CATGAAATGTTGTATCATCACTTGGATCAACACTTTCAATAATTGGAGTAATATTTTTTA  420
Hesp_04    CATGAAATGTTGTATCATCACTTGGATCAACACTTTCAATAATTGGAGTAATATTTTTTA  420
Hesp_02    CATGAAATGTTGTATCATCACTTGGATCAACACTTTCAGTAATTGGAGTAATATTTTTTA  420
           ************************************** *********************
```

Figure S1 page 2/3

**B: *Notonecta* sp.**

```
Noto_05    GGACAGTAATCAACTATAGACCTTCCGTAATATGAGCCCTGGGATTCGTATTTTTATTTA  60
Noto_06    GGACAGTAATCAACTATAGACCTTCCGTAATATGAGCCCTGGGATTCGTATTTTTATTTA  60
Noto_08    GGACAGTAATCAACTATAGACCTTCCGTAATATGAGCCCTGGGATTCGTATTTTTATTTA  60
Noto_07    GGACAGTAATCAACTATAGACCTTCCGTAATATGAGCCCTGGGATTCGTATTTTTATTTA  60
           ************************************************************

Noto_05    CATTAGGGGGATTAACAGGAGTTGTATTAGCTAATTCATCAATTGATATCATTCTACATG  120
Noto_06    CATTAGGGGGACTAACAGGAGTTGTATTAGCTAATTCATCAATTGATATCATTCTACATG  120
Noto_08    CATTAGGGGGACTAACAGGAGTTGTATTAGCTAATTCATCAATTGATATCATTCTACATG  120
Noto_07    CATTAGGGGGACTAACAGGAGTTGTATTAGCTAATTCATCAATTGATATCATTCTACATG  120
           *********** ************************************************

Noto_05    ATACATATTACGTAGTAGCACATTTTCACTATGTATTATCAATAGGAGCTGTATTTGCAA  180
Noto_06    ATACATATTATGTAGTAGCACATTTTCACTATGTATTATCAATAGGAGCTGTATTTGCAA  180
Noto_08    ATACATATTACGTAGTAGCACATTTTCACTATGTATTATCAATAGGAGCTGTATTTGCAA  180
Noto_07    ATACATATTACGTAGTAGCACATTTTCACTATGTATTATCAATAGGAGCTGTATTTGCAA  180
           ********** *************************************************

Noto_05    TCATTGGAAGATTTATTCAATGATACCCCTTATTTACTGGAGTAACAATAAATCCTAAAT  240
Noto_06    TCATTGGAAGATTTATTCAATGATACCCCTTATTTACTGGAGTAACAATAAATCCTAAAT  240
Noto_08    TCATTGGAAGATTTATTCAATGATACCCCTTATTTACTGGAGTAACAATAAATCCTAAAT  240
Noto_07    TCATTGGAAGATTTATTCAATGATACCCCTTATTTACTGGAGTAACAATAAATCCTAAAT  240
           ************************************************************

Noto_05    GATTAAAGATACACTTTATAATTATATTTGTAGGAGTAAATATAACATTTTTTCCTCAAC  300
Noto_06    GATTAAAGATACACTTTATAATTATATTTGTAGGAGTAAATATAACATTTTTTCCTCAAC  300
Noto_08    GATTAAAGATACACTTTATAATTATATTTGTAGGAGTAAATATAACATTTTTTCCTCAAC  300
Noto_07    GATTAAAGATACACTTTATAATTATATTTGTAGGAGTAAATATAACATTTTTTCCTCAAC  300
           ************************************************************

Noto_05    ATTTCTTAGGACTAAGAGGAATACCTCGACGTTATTCTGATTACCCAGATAGATTTACAA  360
Noto_06    ATTTCTTAGGACTAAGAGGAATACCTCGACGTTATTCTGATTACCCAGATAGATTTACAA  360
Noto_08    ATTTCTTAGGACTAAGAGGAATACCTCGACGTTATTCTGATTACCCAGATAGATTTACAA  360
Noto_07    ATTTCTTAGGACTAAGAGGAATACCTCGACGTTATTCTGATTACCCAGATAGATTTACAA  360
           ************************************************************

Noto_05    CATGAAACGTTGTATCATCTATTGGCTCTACAATATCAGTAGTAGGGGTAGCCATATTCA  420
Noto_06    CATGAAACGTTGTATCATCTATTGGCTCTACAATATCAGTAGTAGGGGTAGCCATATTCA  420
Noto_08    CATGAAACGTTGTATCATCTATTGGCTCTACAATATCAGTAGTAGGGGTAGCCATATTCA  420
Noto_07    CATGAAACGTTGTATCATCTATTGGATCTACAATATCAGTAGTAGGGGTAGCCATATTCA  420
           ************************* **********************************

Noto_05    TCTTCATTATATGAGAAAGATTAGTTGCTAAACGGACTATTATATTCCCAAATAATATAA  480
Noto_06    TCTTCATTATATGAGAAAGATTAGTTGCTAAACGGACTATTATATTCCCAAATAATATAA  480
Noto_08    TCTTCATTATATGAGAAAGATTAGTTGCTAAACGGACTATTATATTCCCAAATAATATAA  480
Noto_07    TCTTCATTATATGAGAAAGATTAGTTGCTAAACGGACTATTATATTCCCAAATAATATAA  480
           ************************************************************
```

Figure S1 page 3/3

**C: *Laccophilus macuolosus***

```
Lacc_11    GATCACAAATTAGATATAGACCATCTTTACTTTGAGCATTAGGATTTGTATTTTTATTTA  60
Lacc_09    GATCACAAATTAGATATAGACCATCTTTACTTTGAGCATTAGGATTTGTATTTTTATTTA  60
Lacc_10    GATCACAAATTAGATATAGACCATCTTTACTTTGAGCATTAGGATTTGTATTTTTATTTA  60
           ************************************************************

Lacc_11    CTGTTGGGGGTTTAACTGGAGTAGTATTAGCAAATTCATCAATTGATATTATTCTTCATG  120
Lacc_09    CTGTTGGGGGTTTAACTGGAGTAGTATTAGCAAATTCATCAATTGATATTATTCTTCATG  120
Lacc_10    CTGTTGGGGGTTTAACTGGAGTAGTATTAGCAAATTCATCAATTGATATTATTCTTCATG  120
           ************************************************************

Lacc_11    ATACATATTATGTTGTAGCTCATTTTCACTATGTATTATCTATAGGAGCTGTATTTGCTA  180
Lacc_09    ATACATATTATGTTGTAGCTCATTTTCACTATGTATTATCTATAGGAGCTGTATTTGCTA  180
Lacc_10    ATACATATTATGTTGTAGCTCATTTTCACTATGTATTATCTATAGGAGCTGTATTTGCTA  180
           ************************************************************

Lacc_11    TTCTTGCGGGATTTATTCAATGATTCCCATTATTTACCGGAATCACATTAAATTCTAAAT  240
Lacc_09    TTCTTGCGGGATTTATTCAATGATTCCCATTATTTACCGGAATCACATTAAATTCTAAAT  240
Lacc_10    TTCTTGCGGGATTTATTCAATGATTCCCATTATTTACCGGAATCACATTAAATTCTAAAT  240
           ************************************************************

Lacc_11    TATTAAAAATTCAATTTATTGTAATATTTATTGGAGTAAATTTAACTTTCTTTCCTCAAC  300
Lacc_09    TATTAAAAATTCAATTTATTGTAATATTTATTGGAGTAAATTTAACTTTCTTTCCTCAAC  300
Lacc_10    TATTAAAAATTCAATTTATTGTAATATTTATTGGAGTAAATTTAACTTTCTTTCCTCAAC  300
           ************************************************************

Lacc_11    ATTTCTTAGGATTAAGAGGTATACCTCGTCGTTATTCCGATTACCCAGATGCCTATACAT  360
Lacc_09    ATTTCTTAGGATTAAGAGGTATACCTCGTCGTTATTCCGATTACCCAGATGCCTATACAT  360
Lacc_10    ATTTCTTAGGATTAAGAGGTATACCTCGTCGTTATTCCGATTACCCAGATGCCTATACAT  360
           ************************************************************

Lacc_11    CATGAAATGTAATTTCTTCTATTGGATCGACTATTTCATTTATTGGAGTATTATTATTAA  420
Lacc_09    CATGAAATGTAATTTCTTCTATTGGATCGACTATTTCATTTATTGGAGTATTATTATTAA  420
Lacc_10    CATGAAATGTAATTTCTTCTATTGGATCGACTATTTCATTTATTGGAGTATTATTATTAA  420
           ************************************************************

Lacc_11    TTTATATTATTTGAGAAGCATTTATTTCACAACGAATAGTTATTTTTTCTAATCAAATAC  480
Lacc_09    TTTATATTATTTGAGAGGCATTTATTTCACAACGACTAGTTATTTTTTCTAATCAAATAC  480
Lacc_10    TTTATATTATTTGAGAAGCATTTATTTCACAACGACTAGTTATTTTTTCTAATCAAATAC  480
           **************** ****************** ************************
```

**Figure S1 (above). Alignments of partial *COI* (*cytochrome oxidase subunit I*) from (A) *Hesperocorixa* sp. (Hesp), (B) *Notonecta* sp. (Noto), and (C) *Laccophilus maculosus* (Lacc) collected from two ponds in Eastern Nova Scotia, Canada.** Sequences were aligned using Clustal Omega, and asterisks indicate identical nucleotides among all sequences in the alignment.

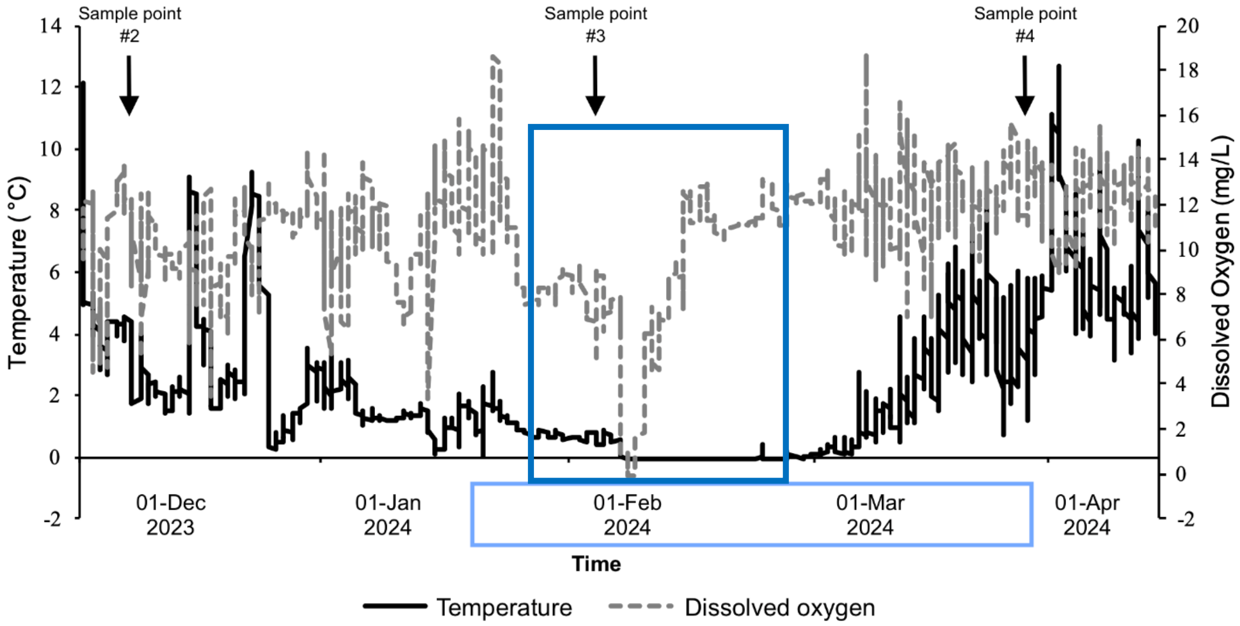


**Figure S2. Temperature and dissolved oxygen concentration recorded in Griffith Pond from October 2023 to April 2024.** The solid, black line represents temperature, and the grey, dashed line represents dissolved oxygen. The dark blue rectangle indicates when the logger became encased in snow and ice approximately between 20 January and 20 February 2024. The light blue rectangle represents the period of time when ice cover was sustained. Downward-pointing arrows indicate experimental sampling points used for mesocosms (#2 and 3) or field sites (#4).

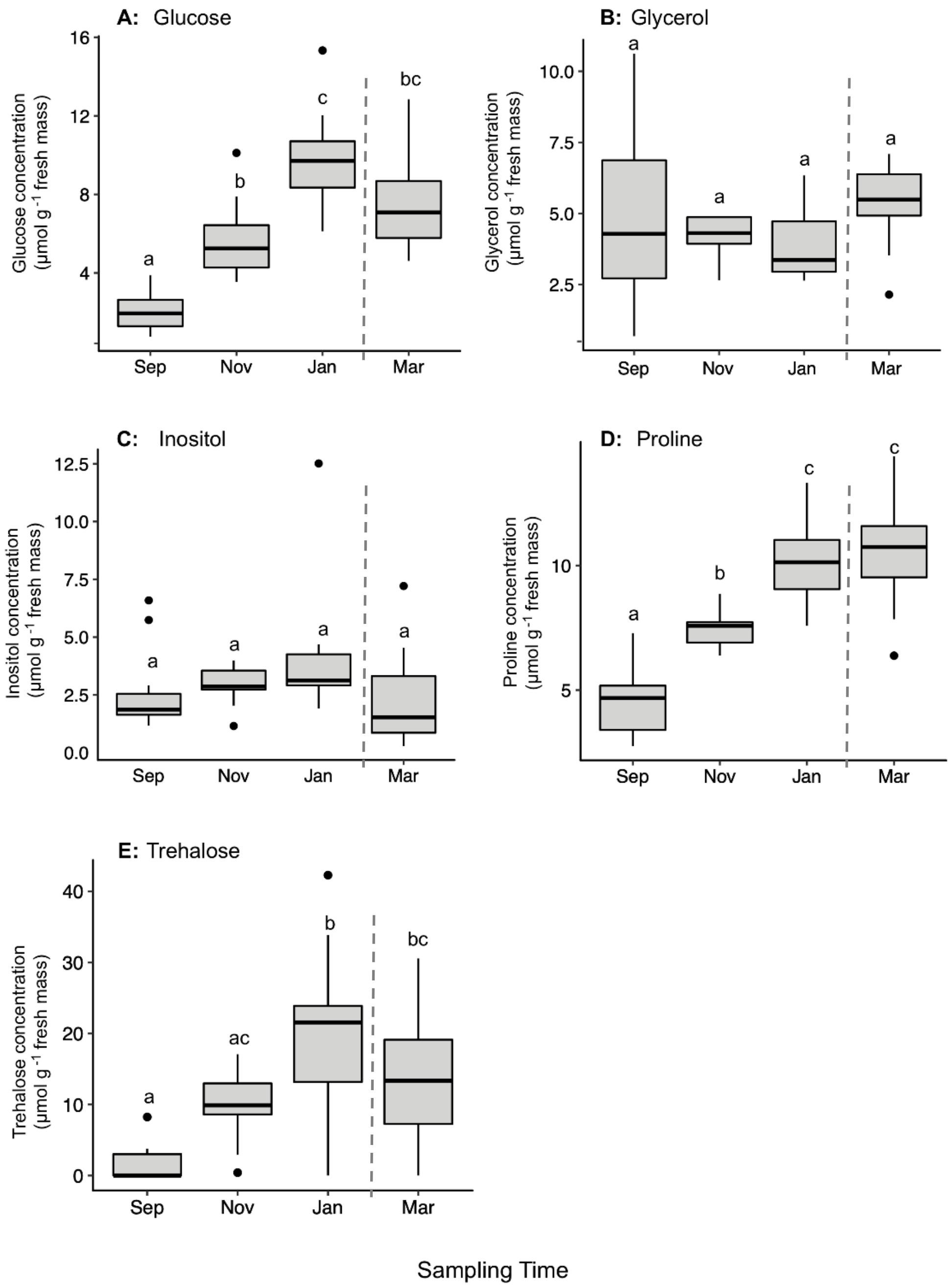


**Figure S3. Spectrophotometrically determined concentration of (A) glucose, (B) glycerol, (C) *myo*-inositol, (D) proline and (E) trehalose in backswimmers (*N* = 50) sampled from mesocosms in September 2023, November 2023, and January 2024 and from the field in March 2024**. The bottom and top of each box represents the lower and upper quartile, respectively; median is represented by the horizontal line; the vertical lines extend to the maximum and minimum values with 1.5 times the inter-quartile range; outliers are indicated by black dots. Different letters above boxes indicate statistical significance between sampling time points, determined using ANOVA and Tukey's post-hoc analyses ($P < 0.05$).

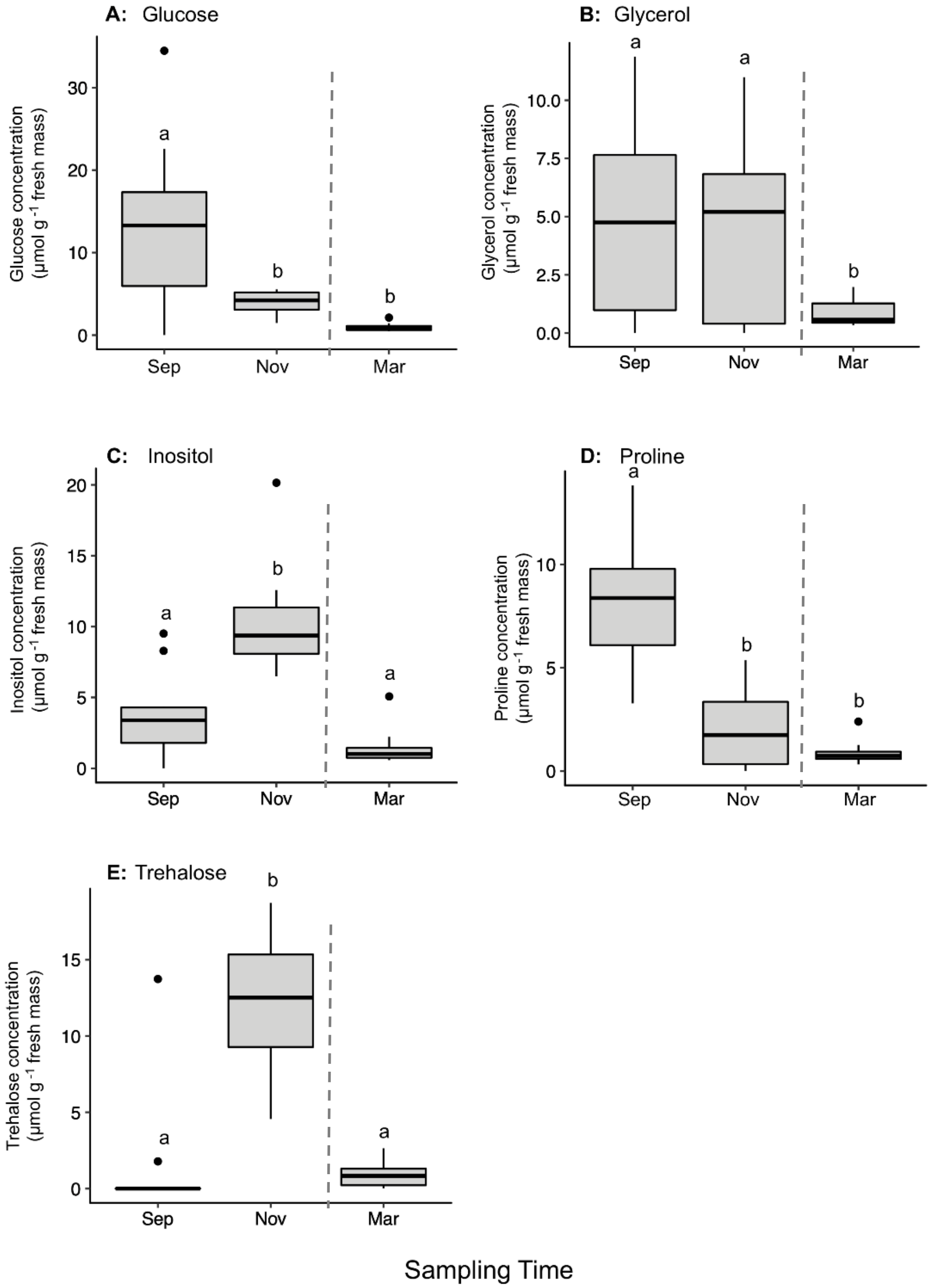


**Figure S4. Spectrophotometrically determined concentration of (A) glucose, (B) glycerol, (C) *myo*-inositol, (D) proline and (E) trehalose in diving beetles (*N* = 36) sampled from mesocosms in September 2023 and November 2023, and from the field in March 2024.** The bottom and top of each box represents the lower and upper quartile, respectively; median is represented by the horizontal line; the vertical lines extend to the maximum and minimum values with 1.5 times the inter-quartile range; outliers are indicated by black dots. Different letters above boxes indicate statistical significance between sampling time points, determined using ANOVA and Tukey's post-hoc analyses ($P < 0.05$).

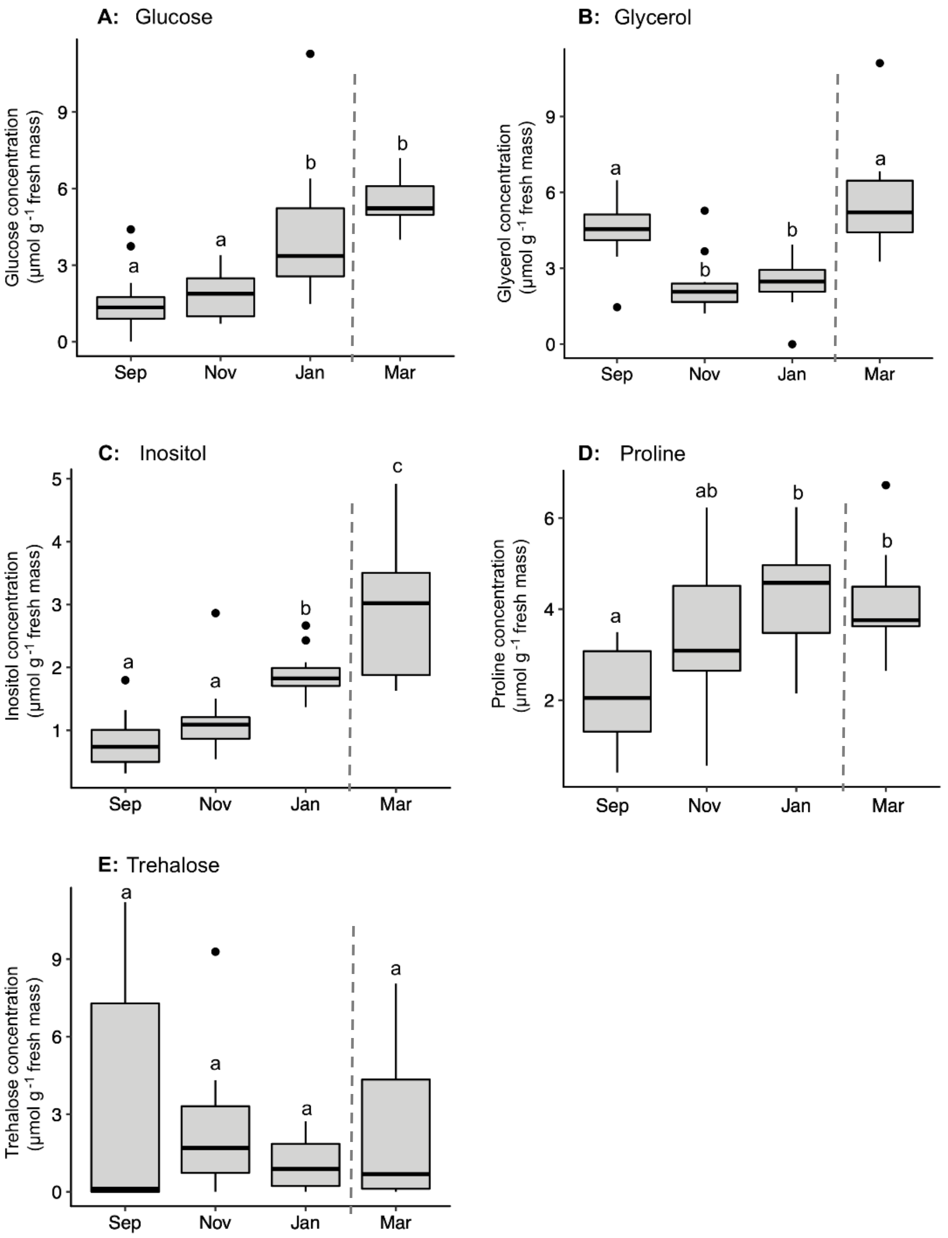


**Figure S5. Spectrophotometrically determined concentration of (A) glucose, (B) glycerol, (C) *myo*-inositol, (D) proline and (E) trehalose in water boatmen (*N* = 52) sampled from mesocosms in September 2023, November 2023, and January 2024 and from the field in March 2024.** The bottom and top of each box represents the lower and upper quartile, respectively; median is represented by the horizontal line; the vertical lines extend to the maximum and minimum values with 1.5 times the inter-quartile range; outliers are indicated by black dots. Different letters above boxes indicate statistical significance between sampling time points, determined using ANOVA and Tukey's post-hoc analyses ($P < 0.05$).
